# A Genome-wide microRNA screen identifies the microRNA-183/96/182 cluster as a modulator of circadian rhythms

**DOI:** 10.1101/2020.09.29.319798

**Authors:** Lili Zhou, Caitlyn Miller, Loren J. Miraglia, Angelica Romero, Ludovic S. Mure, Satchidananda Panda, Steve A. Kay

**Author notes:** Corresponding author: Steve A. Kay, Ph.D., Keck School of Medicine, University of Southern California, Los Angeles, CA 90089, USA. **Author Contributions:** L.Z. and S.A.K. designed research; L.Z., C.M., L.J.M., A.R., and L.S.M. performed search; L.Z. C.M. L.S.M. analyzed data; L.J.M., L.S.M. and S.P. contributed new reagents/ analytic tools; L.Z. wrote the paper.

## Abstract

The regulatory mechanisms of circadian rhythms have been studied primarily at the level of the transcription-translation feedback loops of protein coding genes. Regulatory modules involving non-coding RNAs are less thoroughly understood. In particular, emerging evidence has revealed the important role of miRNAs in maintaining the robustness of the circadian system. To identify miRNAs that have the potential to modulate circadian rhythms, we conducted a genome-wide miRNA screen using U2OS luciferase reporter cells. Among 989 miRNAs in the library, 120 changed the period length in a dosage-dependent manner. We further validated the circadian regulatory function of a miRNA cluster, miR-183/96/182, both *in vitro* and *in vivo*. We found that all three members of this miRNA cluster can modulate circadian rhythms. Particularly, miR-96 directly targeted a core circadian clock gene, PER2. The knockout of the miR-183/96/182 cluster in mice showed tissue-specific effects on circadian parameters and altered circadian rhythms at the behavioral level. This study identified a large number of miRNAs, including the miR-183/96/182 cluster, as circadian modulators. We provide a resource for further understanding the role of miRNAs in the circadian network and highlight the importance of miRNAs as a novel genome-wide layer of circadian clock regulation.

**Significance Statement:** Although miRNAs are emerging as important regulators of diverse physiological and pathological processes, our knowledge of their potential role in regulation of circadian rhythms is still limited. We deployed a cell-based genome-wide screening approach, and successfully identified mature miRNAs as cell-autonomous circadian modulators. We then specifically focused on the miR-183/96/182 cluster among the candidate miRNA hits and revealed their circadian function both *in vitro* and in *vivo* from the unbiased screen. This study provides resources for further understanding the role of miRNAs in the circadian network. It also highlights the importance of miRNAs as a novel genome-wide layer of circadian clock regulation.

## Introduction

Most organisms on earth have internal timing systems that optimally adapt and respond to 24- hour light-dark cycles. In mammals, the endogenous timing systems function and synchronize at multiple levels, from molecular to neural circuits (1). At the molecular level, the circadian clock is generated by a cell-autonomous regulatory feedback loop, where gene activators, such as BMAL1 and CLOCK, turn on the expression of gene repressors, such as *PER* and *CRY*, which in turn repress the expression and transcriptional activity of the activators (2, 3). These simplified steady-state mRNA/protein rhythms actually undergo complicated dynamic regulation by many other processes including RNA splicing, polyadenylation, nuclear export, RNA degradation, and microRNAs (miRNAs) regulation (4, 5).

MiRNAs are a class of small non-coding RNAs. Genes encoding miRNAs can be found in the genome either as single genes or as clusters, i.e. miRNAs are localized in the same genomic region (usually less than 10 kb), transcribed in the same orientation, and highly co-expressed (6–8). These genes are first transcribed in the nucleus as a single primary transcript of miRNAs (pri-miRNAs) and are then cleaved to ∼70-nt precursor miRNA (pre-miRNA) hairpins followed by further maturation in the cytoplasm. The biologically active mature miRNAs are single-stranded and ∼22-nt in length (9, 10). In mammals, mature miRNAs usually bind to sites in the 3’ untranslated region (3’UTR) of their target mRNAs through base-pairing of the seed region, mainly at position 2-7 from the 5’end of the miRNA. This base-paring is generally considered the minimal need to engage a target mRNA (10). Beyond the seed region, the binding between the whole mature miRNA sequence and the target mRNA is not perfectly complementary. These imperfect bindings interfere with the stability or translation of mRNAs (11, 12). They result in a complex network where a single miRNA can concurrently regulate a large number of target genes, and one gene can be regulated by multiple miRNAs (13–15). The regulations between miRNAs and target genes are highly time- and tissue-specific (16–18), and play important roles in diverse physiological and pathophysiological processes (19–23).

Although miRNAs are emerging as major regulators of diverse physiological and pathological processes, particularly in cancer (12), the knowledge of their roles in circadian rhythms is still limited to a few genes. To address this, several studies, including some genome-wide profiling experiments, focused on identifying miRNAs expressed in a circadian manner in various tissues and across different species, including mouse, *Drosophila*, and *Arabidopsis* (24–27). The focus of these studies was clock-controlled miRNAs, but the observed rhythmic expression alone of these miRNAs is neither necessary nor sufficient for them to be considered as regulators of circadian rhythms. Most of these studies did not determine how the oscillation of the miRNAs is relevant to specific physiological and behavioral rhythms. Instead, more convincing evidence for the regulatory role of miRNAs in clock function came from a genetic study in mice, where global knockout of *Dicer*, the enzyme responsible for cleaving pre-miRNA to generate mature miRNAs, significantly shortened the circadian period (28). Several other studies have also investigated specific miRNAs that regulate the expression of core clock components, for example, miR-142 regulates *Bmal1*, miR-17 regulates *Clock,* and miR-192/194 cluster regulates the *Per* gene family (29–32). These studies together suggest a pervasive regulation of the clock by miRNAs that requires further functional and mechanistic understanding.

In 2007, Xu et al. identified a sensory organ-specific miRNA cluster of miR-183, miR-96, and miR-182, named the miR-183/96/182 cluster. Members of the miR-183/96/182 cluster are located within a 4-kb genomic region on mouse chromosome 6qA3, and this genomic arrangement is highly conserved in mammals (27). Some speculation exists regarding a circadian function for the miR-183/96/182 cluster or its members for several reasons. First, the gene expression of the miR-183/96/182 cluster oscillates in mouse retina in a circadian manner (27). Second, the predicted targets of the miR-183/96/182 cluster include some circadian rhythm relevant genes, and Adenylyl cyclase type 6 (*Adcy6*), a clock-controlled gene that modulates melatonin synthesis, is expressed in anti-phase with the miR-183/96/182 cluster (27). Third, in humans, a polymorphism in pre-miR-182 was found to be associated with a marked reduction in the expression of the circadian gene *CLOCK* (33). However, to date, there has been no direct evidence to assign specific roles of the miR-183/96/182 cluster in the modulation of circadian rhythms.

In the present study, we used a cell-based genome-wide screening approach to systematically identify the mature miRNAs that can potentially alter circadian rhythms. We show here that our cell-based miRNA screening successfully identified cell-autonomous circadian modulators. We specifically focused on the miR-183/96/182 cluster among the candidate miRNA hits and revealed their circadian function both *in vitro* and in *vivo* from the unbiased screen.

## Results

### 2.1 Genome-wide miRNA screen

To identify miRNAs that have the potential to modulate circadian rhythms, we first adopted a mammalian cell-based assay to screen a library of miRNAs (Figure 1A). We used U2OS cells which contain either the *Bmal1-dLuc* or *Per2-dLuc* gene to report circadian oscillations (34). We screened a Qiagen miRNA library containing 989 synthesized human miRNA mimics, which are chemically synthesized small RNA molecules imitating mature miRNAs. An equal amount of these miRNA mimics was pre-spotted in each well of 384-well plates individually, and then reverse transfected into cells. By integrating the screening results from both *Bmal1-dLuc* and *Per2-dLuc* reporter cell lines, we found 113 long-period hits (11%) and 7 short-period hits (1%) from both sets of screens (Figure 1B, Table S1). In addition, we observed many hits that only show low amplitude rather than period change. Considering the low amplitude phenotypes were most likely caused by non-circadian related cell fitness issues rather than circadian clock deficiency, these miRNA hits were not included in the follow-up studies.

**Figure 1.**
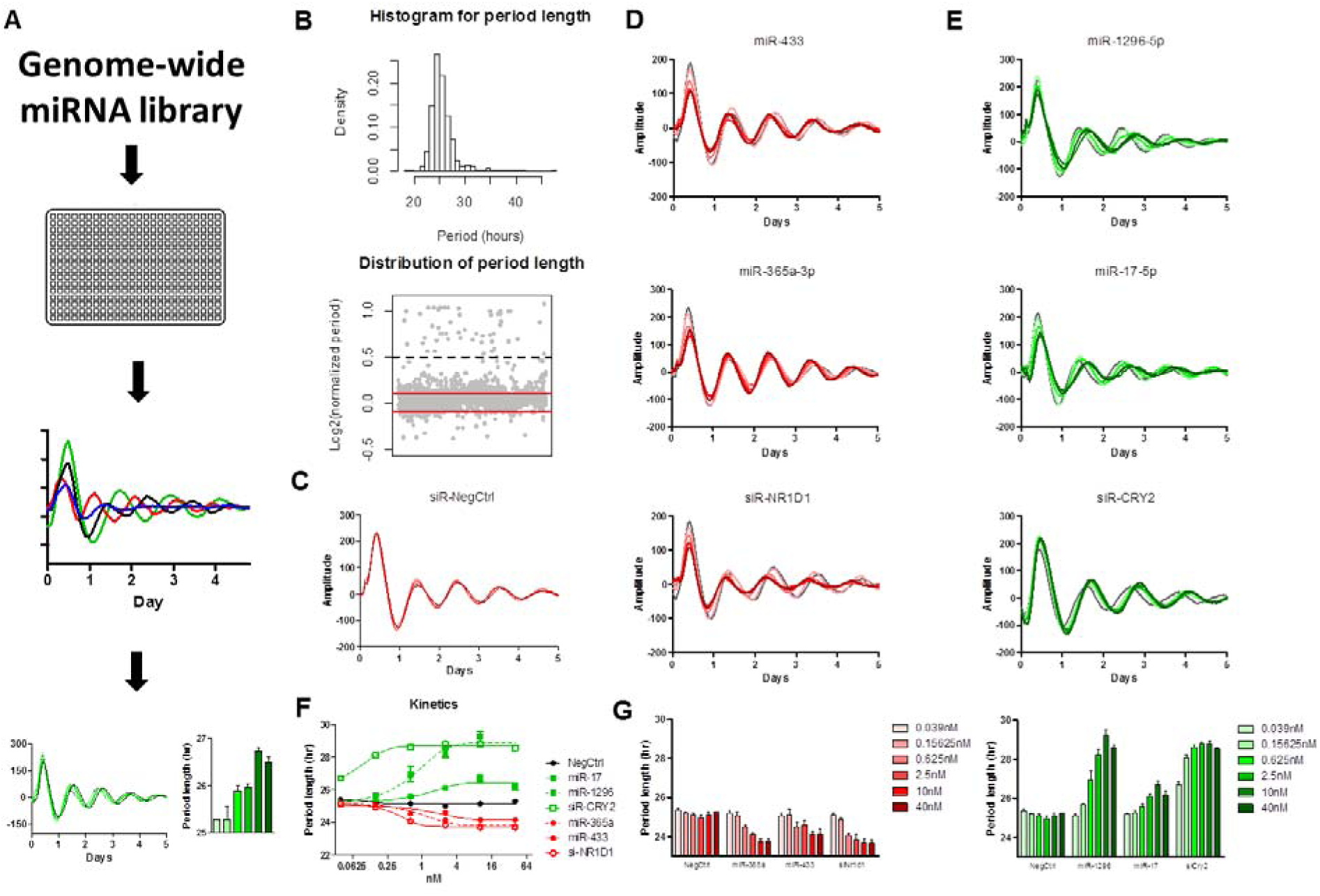
A cell-based genome-wide screen for miRNA modifiers of circadian rhythms. (A) A schematic diagram of the genome-wide miRNA screen. *Bmal1-dLuc* and *Per2-dLuc* reporter cells were transfected with miRNAs in 384-well plates. Kinetic bioluminescence was recorded and analyzed to obtain circadian parameters and select candidate hits. Validation was performed by measuring the dose-dependent effects of some candidate miRNAs on circadian phenotypes. (B) Distribution of circadian period lengths of the screen. The histogram shows a tail torward the long period. Gray dots in the botom panel represent normalized period lengths. The period length fo each well was divided by the negative control and indicated in Log2 space. The cut-off was - 0.1 and +0.1 (red lines). Values above 0.5 (black dash line) were considered outlier. (C) Representative bioluminescence profiles for dose-dependent phenotypic validation of negative control, short period positive control (siNR1D1), long period positive control (siCRY2). (D) short period candidate hits (miR-433, miR-365a-3p) and (E) long period candidate hits (miR-1296-5p, miR-17). (F) The kinetics of dose-dependent effects of the indicated candidate miRNAs and control RNAs. (G) The period length parameters of the indicated candidate miRNAs and control RNAs. *Per2-dLuc* U2OS cells were transfected with the indicated amount of miRNAs (six 4-fold dilution series from 0.039 to 40 nM/well). Data represents the mean ± SD (n = 4).

Based on sequence similarity in the seed regions (35), miRNAs can be classified into families, which are usually evolutionary related but do not strictly share common ancestry (10). The similar seed regions between members in the same family, in theory, lead to similar functions. Among the 120 candidate hits, 65 (54%) miRNAs belong to 35 families that have more than two members in humans (e.g. multi-member family), and the rest of the hits are in families that currently contain only one member, themselves (Table S2). Some of these identified families were highly enriched in candidate hits. For example, eight miRNAs from family let-7 and four miRNAs from family miR-17 were identified and showed a circadian phenotype in the screen. These might be attributed to similar seed regions shared by members of the miRNA family.

The other way to group miRNAs is based on their genomic locations because miRNAs tend to co-localize as clusters (6). Using all human miRNA data from miRBase (35), miRNAs are clustered if two neighboring miRNAs are on the same strand and within 10 kb, and in human approximately 20% of all miRNAs are clustered (36). In our screen, 54 out of the 120 miRNAs hits identified were mapped to 35 clusters (Table S3). This number accounts for 45% of all miRNAs hits identified in the screen, which is higher than the overall ratio in all human miRNAs. Although miRNA clusters are defined by physical distance rather than the sequence similarity, we noticed that some clusters were enriched in candidate hits. For instance, all three members in cluster let-7a-1/let-7f-1/let-7d and all three members in cluster miR-183/miR-96/miR-182 were identified by the screen. Based on the seed sequence similarity, we further divided the 35 identified clusters into three different categories (36): 7 homo-seed clusters (miRNA members having identical seed sequences), 19 hetero-seed clusters (miRNA members having different seed sequences), and 9 homo-hetero-seed clusters (a combination of the former two classes). We found that homo-seed clusters have a significantly higher percentage of candidate miRNA enrichment than that of hetero-seed clusters and homo-hetero-seed clusters (Figure S1).

We then randomly selected some candidate miRNA hits and conducted a dose-dependent assay for further validation. We tested these miRNAs at 6 different doses (4-fold dilution series; from 40 to 0.039 nM). All of them showed dose-dependent effects on circadian parameters (Figure 1C-E). The kinetics of miRNAs dose-dependent effects showed that the phenotype reaches a plateau at about 10 nM, which was the dosage used in our miRNA screen (Figure 1F, G). In addition to miRNA, small-interfering RNA (siRNA) is another class of small RNAs (synthetic double-strand RNA, ∼20-25 bp in length) that is central to RNA interference (37). Despite differences in origins, siRNAs and miRNAs share similarities in terms of downstream cellular machinery (37). We, therefore, compared the dose-dependency of miRNAs with that of siRNAs. We found that the potency of the miRNAs was generally lower than that of the siRNAs in our dose-dependent assay (Figure 1D-G). For example, the CRY2 siRNA could significantly shorten the circadian period length of U2OS-reporter cells at a concentration lower than 0.02 nM, but the miRNAs typically did not show a circadian phenotype in reporter cells at a concentration lower than 0.5 nM.

### 2.2 The miR-183/96/182 cluster was important for maintaining circadian period length and amplitude in human cells

The miR-183/96/182 cluster drew our attention because all members from this cluster were identified by the screen. This cluster is conserved in both human and mouse, and has members sharing similar though not identical seed regions (Figure 2A-B). This miRNA cluster was suggested to be involved in circadian rhythms given that the predicted targets include some circadian relevant genes (33, 38). However, no direct evidence has validated the role of this miRNA cluster in circadian rhythm modulation. In our study, all members from this cluster showed a longer period and lower amplitude phenotype after being overexpressed in reporter cells, in a dose-dependent manner (Figure 2C-D).

**Figure 2.**
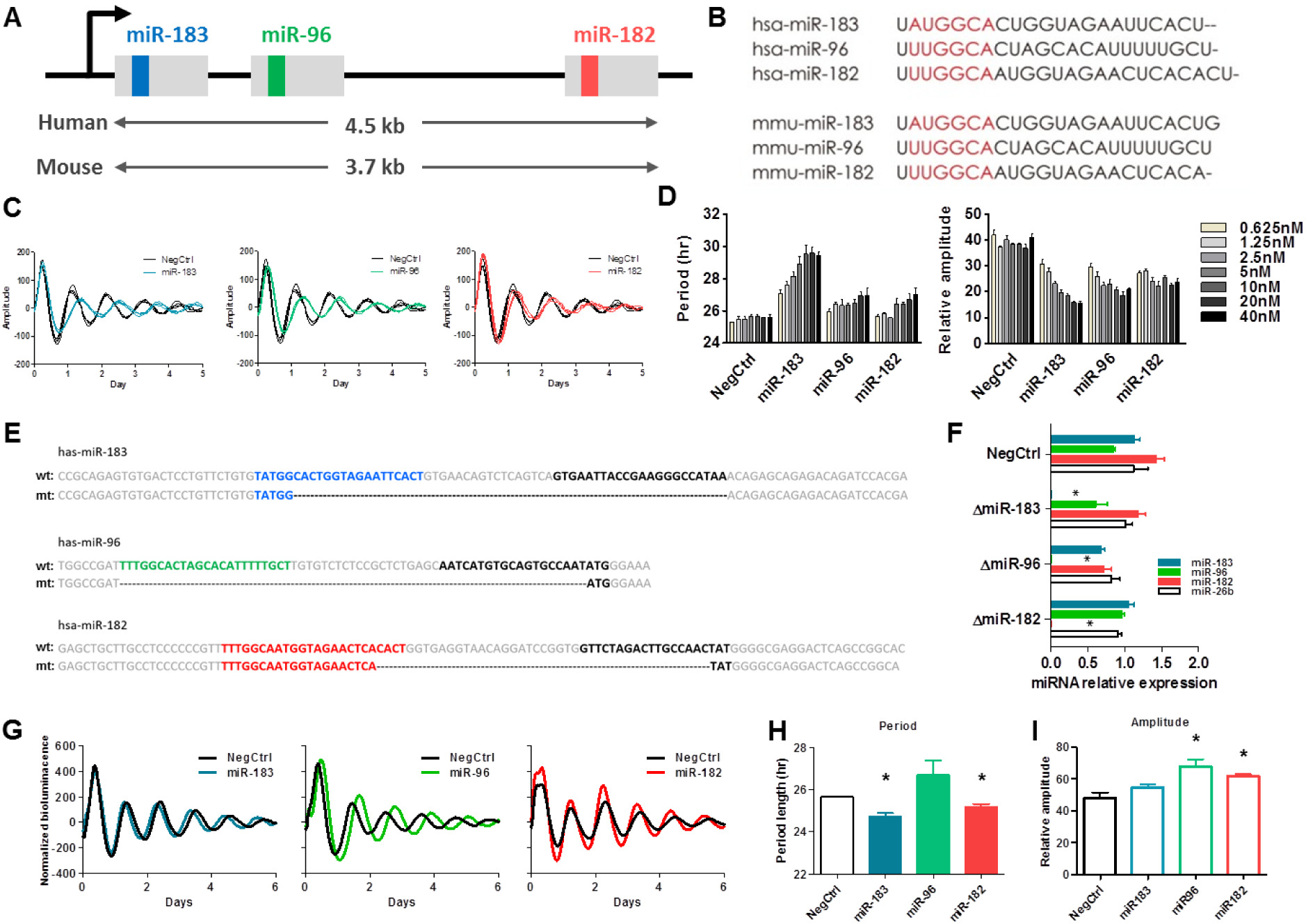
The cluster of miR-183/96/182 altered the circadian period length and amplitude at the cellular level. (A) Schematic diagram showing the miR-183/96/182 cluster on the chromosome. Members of the miR-183/96/182 cluster locate on the same chromosome, and are co-transcribed through the same promoter. The gray blocks represent the primary miRNA. The inside color boxes (blue, green and red) represent the sequences for miR-183-5p, miR-96-5p and miR-182-5p respectively. (B) The sequences of mature miRNA for members of the miR-183/96/182 cluster in human and mouse. The red region represents the seed region of each miRNA, which exhibits high similarity. (C) The dose-dependent effects of members of the miR-183-/96/182 cluster on period length. U2OS *Per2-dLuc* reporter cells were transfected with the indicated amount of miRNAs (Five 2-fold dilution series from 2.5 to 40 nM/well). The data represents the mean ± SD (n = 4). (D) Representative bioluminescence profiles of U2OS *Per2-dLuc* reporter cells which were transfected with 10nM synthetic miRNA mimics of each member of the miR-183/96/182. (E) Knock-out of each member of the miR-183/96/182 cluster using a CRISPR-Cas9 approach. For each miRNA, the “wt” sequence represents the sequence of wild-type primary miRNA, and the “mt” sequence represents the mutant sequence of primary miRNA after deletion. The colored sequences (blue, green and red) represent the 5’miRNA, and the black sequences represent the 3’ miRNA. Deletions are marked by dashes. (F) The mature miRNA expression levels detected by qPCR confirmed the complete deletion of each miRNA. (G) Representative bioluminescence profiles and period lengths for each CRISPR miRNA deletion cell line. Data represent the mean ± SEM (n = 3), * *p* < 0.05.

To confirm that the observed circadian phenotypes were not due to ectopic effects generated by the massive transfection of synthetic miRNAs, we generated miRNA knockout reporter cell lines using a CRISPR-Cas9 approach. Because of their very short length and non-coding properties, there is very limited space in miRNAs for identifying protospacer adjacent motif (PAM) sequences. Small indels induced by CRISPR-Cas9 do not cause a frameshift in the non-coding gene, and may be tolerable during the imperfect complementary binding between miRNAs and their targets. We therefore chose to use paired CRISPR RNAs (crRNAs) to individually excise a genomic sequence encoding each member of the miR-183/96/182 cluster (Figure 2E), resulting in a complete loss of expression of the mature miRNAs (Figure 2F). The knockout of miR-183 or miR-182 shortened the circadian period length (Figure 2G-H), and the knockout of miR-96 or miR-182 increased the amplitude (Figure 2I). These results were opposite from the observed effects when the miRNA mimics were transfected in the cells. These results indicate an intrinsic effect of the miR-183/96/182 cluster on circadian rhythms.

### 2.3 Circadian behaviors were altered in *miR-183/96/182* cluster deficient mice

Next, we analyzed the circadian behaviors of *miR-183/96/182* cluster deficient mice in running-wheel cages. The *miR-183/96/182* cluster gene-trap (*miR-183C^GT/GT^*) mouse has the expression of the *miR-183/96/182* cluster inactivated throughout the whole body (38). Compared with wild-type littermate control mice, heterozygous *miR-183C^GT/+^* mice had shorter free-running periods in constant darkness, and homozygous *miR-183C^GT/GT^* mice became arrhythmic after release into constant darkness (Figure 3A-C). Voluntary wheel-running activity is a subset of the total general activity. Access to running wheels may artificially affect the activity patterns. We noticed that the wheel-running activity level was extremely low in *miR-183C^GT/GT^* mice (Figure S2A), although the *miR-183C^GT/GT^* mice were hyperactive compared with wild-type mice (Figure S2B). We found that the *miR-183C^GT/GT^* mice had circling behaviors, which might be caused by vestibular dysfunction (39). The discrepancy between wheel-running activity and total general activity in *miR-183C^GT/GT^* mice suggested reduced access to running wheels. The reduced wheel-running activity might flatten the diurnal rhythmicity (40), causing a false-positive scoring of arrhythmicity. To preclude such possibility, we compared the non-wheel beam break total activities between *miR-183C^+/+^* and *miR-183C^GT/GT^* mice. About one week after release into constant darkness, compared with wild-type controls, the *miR-183C^GT/GT^* mice gradually lost their rhythmicity in general total activity (Figure 3F, G). Furthermore, the *miR-183C^GT/GT^* mice also exhibited a phase advance compared with wild-type mice under entrained conditions (Figure 3H). It has been found that the deficiency of the *miR-183/96/182* cluster can cause retinal degeneration (38, 41–43). As intrinsically photosensitive retinal ganglion cells (ipRGCs) in the retina are the main conduit of light information to the master circadian clock, the suprachiasmatic nucleus (SCN) (44, 45), we asked if the circadian disruption in the *miR-183C^GT/GT^* mice may be caused by dysfunction of light entrainment. We found that the sensitivity of ipRGCs, tested using the pupillary light reflex (PLR), was not significantly different between *miR-183C^+/+^* and *miR-183C^GT/GT^* mice and therefore excluded this possibility (Figure S3). Together, these results validated a functional role of the *miR-183/96/182* cluster in circadian regulation at the behavioral level.

**Figure 3.**
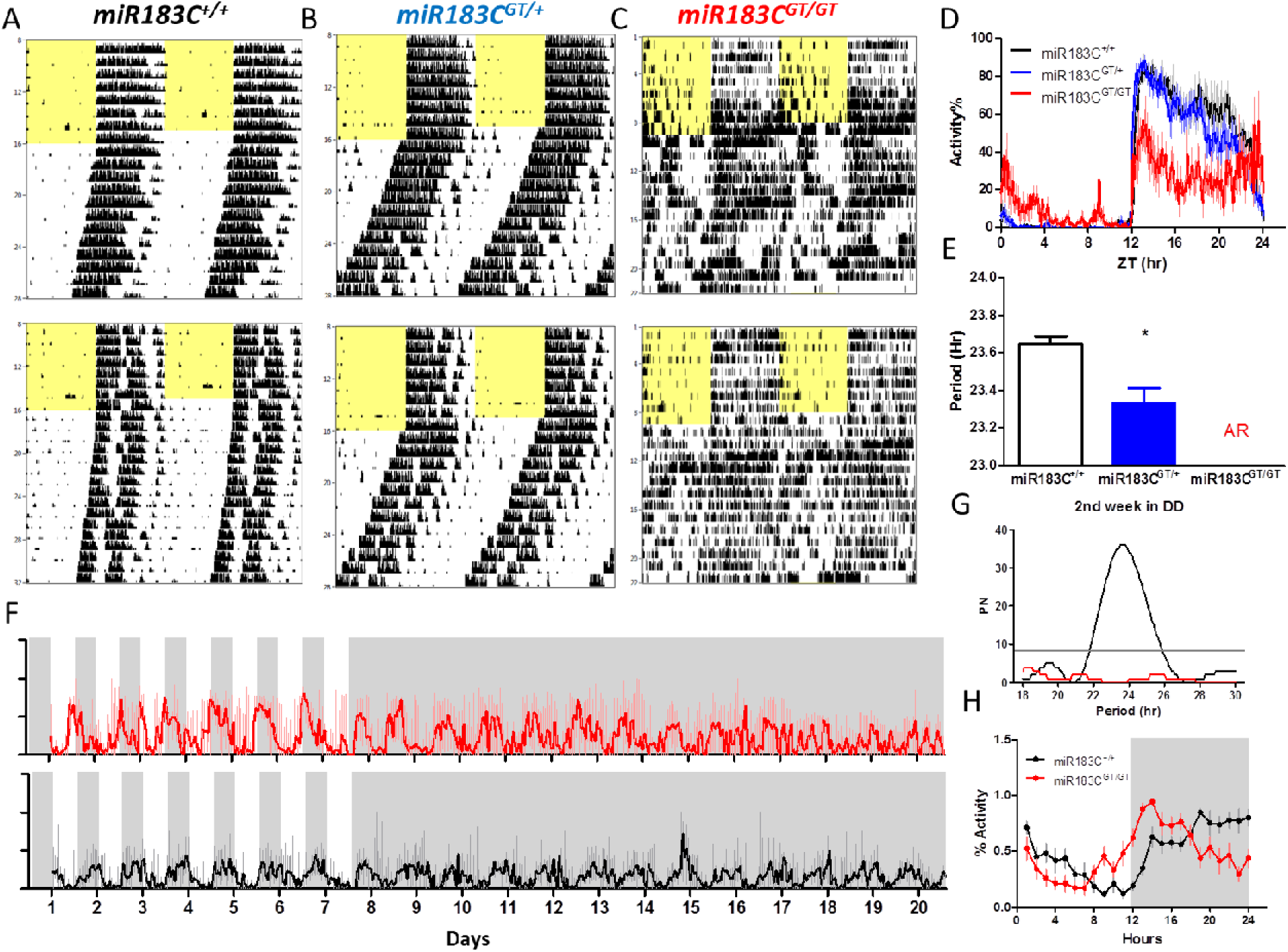
Circadian behaviors were altered in the *miR-183/96/182* cluster deficient mice. (A-C) Representative wheel-running locomotor activity profiles of homozygous wild-type, heterozygous, and homozygous gene trap mice (*miR-183C^+/+^*, n = 14; *miR-183C^GT/+^*, n = 22; *miR-183C^GT/GT^*, n = 10). Animals were maintained on LD12:12 entrained conditions for the first week, indicated by the yellow filled and open area in the records, and then were released to DD for three weeks to measure free-running periods. (D) Diurnal wheel-running activity profiles under the entrained condition. Wheel-running activities were normalized to its highest activity level of each animal and then averaged across 7 days prior to DD. (E) Free-running periods under constant darkness. The free-running period of heterozygous mice was significantly shorter than that of wild-type mice (t-test: * *p* = 0.004). The *miR-183C^GT/GT^* mice became arrhythmic after released into DD. (F) Beam-break total activities of homozygous wild-type and gene-trap mice (*miR-183C^+/+^*, n = 4; *miR-183C^GT/GT^*, n = 4). Animals were maintained on LD12:12 for the first 7 days, indicated by the filled and empty bars above the records, and then released to DD for 14 days. (G) Diurnal beam-break total activity profiles under the entrained condition. Activities were normalized to the highest activity level of each animal and then averaged across 7 days prior to DD. (H) Circadian periodicity of the beam-break total activity under DD condition was detected by Lomb-Scargle Periodogram. Activity of *miR-183C^GT/GT^* under DD did not pass the threshold for period detection. Data represent the mean ± SD.

### 2.4 The *miR-183/96/182* cluster altered circadian rhythms in a tissue-specific way

The expression of miRNAs is highly tissue-specific (16–18). The *miR-183/96/182* cluster has very high expression in many sensory organs and is considered to be a neuronal specific miRNA cluster (39, 46). To delineate the circadian function of the *miR-183/96/182* cluster at a tissue level, we generated a *miR-183/96/182* cluster luciferase reporter mouse line by crossing a *miR-183C^GT/GT^* mouse with a *Per2::Luc* reporter knock-in mouse (47). The *Per2::Luc* reporter knock-in mouse has a luciferase coding sequence inserted before the endogenous *Per2* stop codon, thus fused with an intact endogenous *Per2*-3’UTR. Mice expressing PER2::LUC fusion protein have indistinguishable behavioral and molecular characteristics compared with wild-type mice (47, 48). We compared *ex vivo* circadian bioluminescence rhythms between wild type and mutant mice in three selected tissues, which have detectable expression of the *miR-183/96/182* cluster: the retina which has high expression level of the cluster, the lung tissue which contains peripheral clock function, and the SCN which contains the master clock in the brain. Interestingly, the three tissues showed different circadian phenotypes (Figure 4). The inactivation of the *miR-183/96/182* cluster shortened the circadian period in the retina (Figure 4 A-B) but did not change period lengths in the lung tissue (Figure 4 E-F) and the SCN (Figure 4 I-J). The phases of all tissues tested were significantly changed by the inactivation of the *miR-183/96/182* cluster, but in different directions. Retina and lung had a phase delay (Figure 4 D-L); whereas SCN had a phase advance (Figure 4H), which was consistant with the phase advance in general activity (Figure 3H). This indicates that the peripherals were not synchronized with the central pacemaker. Additionally, the amplitude of the PER2::LUC fusion protein was significantly higher in the mutant lung tissue than the wild-type lung tissue (Figure 4 K).

**Figure 4.**
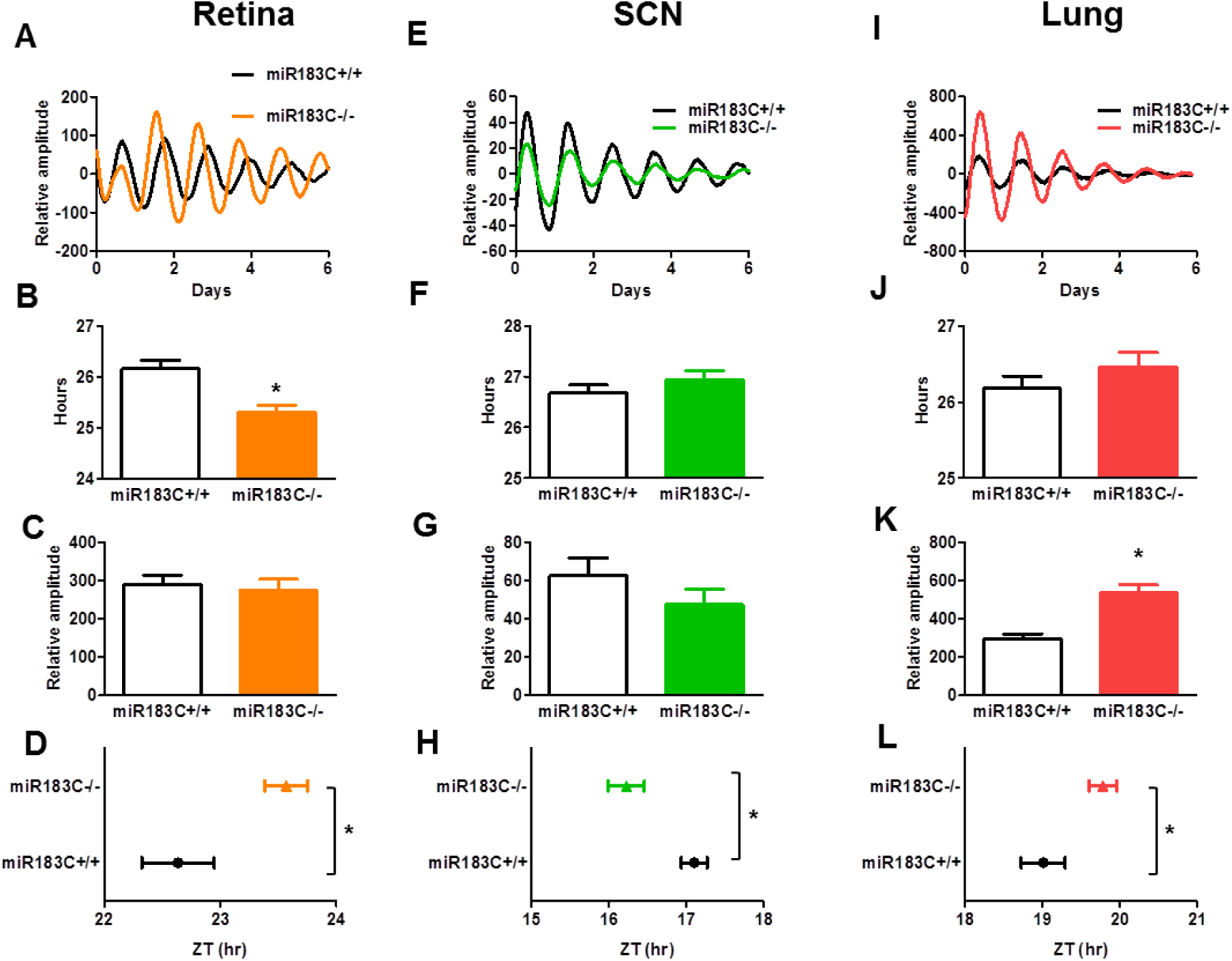
The circadian phenotype of the *miR-183/96/182* cluster was tissue-specific. The *miR-183/96/182* cluster altered the circadian period length, amplitude and phase in the retina (A-D), SCN (E-H), and lung (I-L) from *Per2::Luc* mice carrying homozygous wild-type or homozygous gene-trap alleles (*miR-183C^+/+^*, n = 14∼23; *miR-183C^GT/GT^*, n=12∼19). Animals were maintained under a LD12:12 entrained condition before dissection. Tissue dissection was performed 2-3 hours before lights off and the tissue was immediately cultured for recording for one week. Representative bioluminescence recording profiles between *miR-183C^+/+^* and *miR-183C^GT/GT^* are on the top panel (A, E, and I). Circadian period length (B, F, and J) and relative amplitude (C, G, and K) were calculated by the LM fit (damped sin) method. The phase of each tissue (D, H, and L) was represented by the time of peak luminescence on the first day of constant conditions (first day in Lumicycle), * *p* < 0.05. Data represent the mean ± SD.

### 2.5 PER2 was a direct target of miRNA-96

We further investigated how the miR-183/96/182 cluster contributed to circadian clock regulation and whether there were any core circadian genes among their direct targets. We first used two independent computational methods, DIANA-microT-CDS and miRanda (14, 49, 50), to predict the potential targets for members of the miR-183/96/182 cluster. Among the canonical circadian genes, PER2 and CLOCK were predicted to be direct targets by both methods (Table S4), so we focused on these two genes. Both methods identified the same potential binding region on PER2-3’UTR for miR-96 (Figure 5A). We then validated the direct bindings between miR-96 and PER2-3’UTR by performing a luciferase reporter assay (Figure 5B). Consistently, the over-expression of miR-96 decreased while the knockout of miR-96 increased *PER2* mRNA levels (Figure 5C-D). The over-expression of miR-96 also decreased PER2 at the protein level (Figure 5E-F). Although CLOCK was predicted by both methods to be a target for miR-96, this was not validated by the 3’UTR luciferase reporter assay (data not shown). On the other hand, CLOCK was computationally predicted to be a direct target of miR-182. This was validated through a 3’UTR luciferase reporter assay from other studies (33, 51), as well as in our present study (Figure S4A-B). The over-expression of miR-182, however, did not repress the expression level of CLOCK at either mRNA or protein levels (Figure S4C-D). Due to the complexity of circadian networks, although CLOCK could be directly targeted by the miR-182, the down-regulation effect might be compensated by other factors.

**Figure 5.**
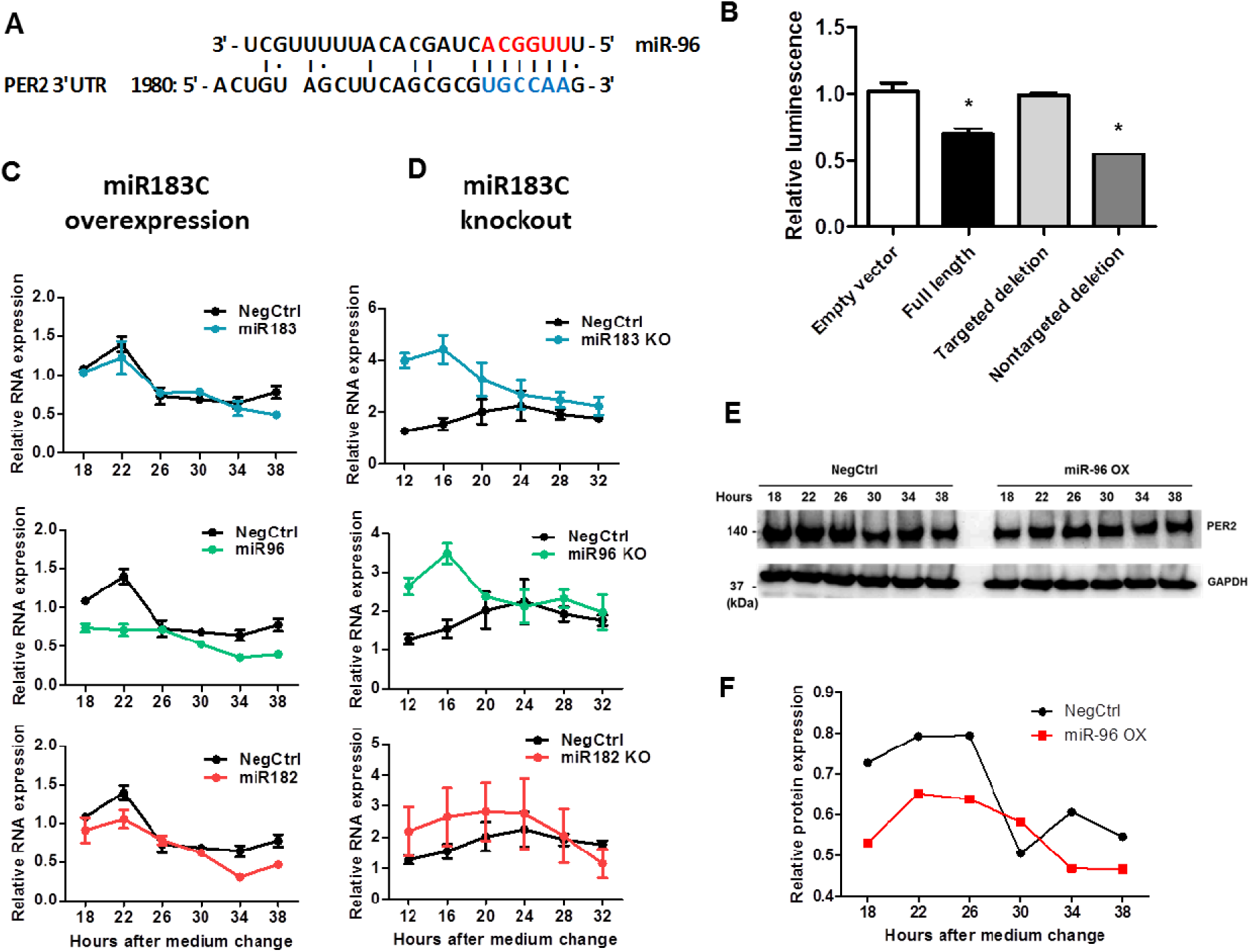
PER2 was a direct target of miR-96. (A) Sequence alignment between miR-96 and its putative binding sites (blue letters) in the PER2-3’UTR. Red letters indicate canonical seed nucleotides. (B) HEK-293T cells were transfected with a luciferase reporter plasmid containing one of the following sequencs: 1) no 3’UTR; 2) a full length wild-type PER2-3’UTR; 3) a full length PER2-3’UTR with a targeted deletion at the predicted binding site of miR-96 (1992_2002 del: gcgcgTGCCAA; upper case is the potential seed binding site of miR-96); 4) a full length PER2-3’UTR with a deletion in a sequence (1712_1734 del: ttcataaacacaagaacacttta) predicted not to be bound by miR-96, * *p* < 0.05. (C) *PER2* mRNA levels in U2OS cells that transfected with miRNA mimics of each member of the miR-183/96/182 cluster. RNA was sampled 18 hours after medium change, every 4 hours across one day. The mRNA levels were determined by qPCR (n=3). (D) *PER2* mRNA levels in miR-183/96/182 cluster knockout cells. RNA from U2OS cells with the deletion of each member of the miR-183/96/182 cluster was harvested 12 hours after medium change, every 4 hours across one day. The mRNA levels were determined by qPCR (n=3). Data represent the mean ± SEM. (E) U2OS cells were transfected with either control or miR-96 mimics, and PER2 protein levels were analyzed by western blot 48 hours post-transfection. (F) Quantification of PER2 protein band intensity compared with the negative control on western blot.

Given the significant phenotypes in circadian rhythms in miR-183/96/182 cluster mutants, we further investigated the expression of some core circadian genes from different mouse tissues as shown in Figure S5. Under light entrained conditions, the gene expression levels of core circadian genes were not changed in all three tissues tested. The peak expression of *Clock*, *Per1*, and *Per2* in mutants was slightly delayed compared with wild types, except for *Per2* that was advanced in the SCN. Under conditions of constant darkness, however, the core circadian genes in the eyes of mutant mice were all significantly lower than those of the wild-type mice. The expression peaks of *Bmal1*, *Clock*, and *Per1* in SCN of mutant mice were advanced compared with those of wild types.

## Discussion

For more than a decade, miRNAs have been found to play important roles in many physiological processes and disease conditions. Although a few miRNAs have been identified as circadian modulators, knowledge of the genome-wide picture of miRNA modulation of circadian rhythms is still limited. Previous studies usually started from either miRNA expression profiling or in silico miRNA target prediction to narrow down specific candidate miRNAs before functionally validating the role of the miRNA as a circadian modulator. In this phenotype-driven study, we started from a genome-wide cell-based functional screen and identified miRNAs that modulate circadian rhythms. The majority of the hits lengthened the period but a very small number of hits were found to shorten the period, which is similar to what was observed in genome-wide siRNA studies (52). Approximately half of the candidate hits could be assigned to a miRNA family or a cluster. Multiple members from a family were likely to be co-identified from the screen, and homo-seed clusters had a higher percentage of the candidate hits, suggesting that the conserved seed sequences were important determinants of functions. However, there were still members from the same family that did not show the same phenotypes. This suggests that differences in secondary structures (53) or in sequences beyond the seed regions still exist in members of the same family, which might be the reason for identifying only a portion in any particular family.

Between the threshold level and saturation level of miRNA concentration, these candidate miRNAs affect the circadian parameters in a dose-dependent manner, but not as efficiently as siRNAs. This may be due to the partial complementary binding between the mammalian miRNA seed regions and their mRNA targets, whereas siRNAs have a perfect binding sequence complementarity to their targets. Ideally, 100 molecues of an siRNA may specificly target 100 molecues of one circadian gene mRNA. However, 100 molecues of an miRNA may target mRNAs of 10 different genes with an average of 10 molecues of each gene. If only one mRNA affects circadian rhythms among the 10 different miRNA target mRNAs, the effect of miRNA on circadian rhtyhms would be greatly diluted. The relatively weaker repression effects of miRNAs compared to that of siRNAs may have their biological implication in the cell. Like a buffer solution requires a mixture of a weak acid and its conjugate base or vice versa, the cellular transcriptome may require a mixture of moderate affinity miRNAs and their conjugate targets for stabilization.

Our screen also identified some miRNA hits that have been previously reported to play a role in circadian rhythms in other studies, which validates our screen as an effective strategy to identify miRNAs as novel circadian clock regulators. For example, 5 out of 6 of the members in the miR-17 cluster were identified from our screen, and *miR-17* was reported to participate in circadian period regulation by directly targeting the *Clock* gene (32). In our screen, we found that the transfection of miR-17 mimics lengthened the circadian period in U2OS reporter cells. Other candidate miRNA hits identified from our screen need further validation for their intrinsic function in circadian modulation. Of course, our screen could not identify all miRNA circadian modulators. One reason is that the miRNA library used in this screen only covers a portion of the currently known miRNA space. A larger size library would be expected to identify more miRNA hits. The phenotype of a given miRNA depends highly depend on the cellular context. Using different reporter cell lines may harvest distinct miRNA hits, given that miRNAs are highly cell-type specific.

Our study focused on the miR-183/96/182 cluster, which has been predicted to be involved in circadian rhythms because of previously observed circadian expression patterns, and the predicted targets include certain core circadian genes or circadian rhythm relevant genes (27, 33, 38). Previous reports link the miR-183/96/182 cluster to circadian rhythms because *Adcy6* is predicted to be a target of *miR-182* and *miR-96*, and *Adcy6* regulates arylalkylamine-N-acetyltransferase (*Aanat*), an enzyme in the melatonin synthesis pathway (27). Melatonin is a hormone produced primarily in the pineal gland and is implicated in the regulation of circadian rhythms in most animals (54). However, most laboratory mouse strains are melatonin deficient (55–58). Therefore, it is less likely that the *miR-183/96/182* cluster affects the circadian rhythms in melatonin deficient mice through modulating the melatonin pathway, although *Adcy6* was experimentally validated to be a target of the *miR-182* and *miR-96* (27). So far, there have been no experiments that validate the function of the miR-183/96/182 cluster as circadian modulators or dissect the circadian role of each member. In the present study, we first determined that each member in this cluster could change the circadian rhythms in human cells through transient transfection of these miRNA mimics. We then confirmed that these circadian effects were endogenous and cell-autonomous by knocking them out in human cells. We found that each member in the miR-183/96/182 cluster exhibits different circadian phenotypes in human cells after being knocked out. Because the seed regions of each member of the miR-183/96/182 cluster are similar but not identical, they have both common and unique targets. Thus, they can show cooperative or opposing effects on biological processes, implying the complexity of this miRNA cluster in mediating gene regulation. We found that PER2 is one of the direct targets of miR-96. However, the validation of CLOCK as a direct target of miR-182 was not conclusive. The expression of endogenous CLOCK was not affected by miR-182, even though the 3’UTR luciferase reporter assay supported the effects. This is probably because the endogenous CLOCK expression is affected by other factors in a complex network context that is not limited to miR-182.

The expression of miRNAs has been shown to be highly tissue-specific (16–18). Consequently, the same miRNAs may behave differently in different tissues. This was observed in our study. Using *Per2::Luc* circadian reporter knock-in mice, we found that different tissues exhibited different circadian phenotypes. In *miR-183/96/182* cluster deficient *Per2::Luc* mice, the SCN showed phase advance, the lung had both phase delay and increased amplitude, and the retina showed a phase delay and shortened period compared with wild-type mice. These differences may be due to the tissue-specific gene expression of the miRNAs themselves, the tissue-specific gene network context, or the interactions between both. The SCN is believed to play an important role in synchronizing the phases of peripheral clocks throughout the body, however, this was not observed in the *miR-183/96/182* cluster deficient mice. We noticed that the phases of the lung and the retina were not synchronized by the SCN in the *miR-183C^GT/GT^ Per2::Luc* mice. The advanced SCN did not move the lung and retina phases forward accordingly in mutant mice. The reason for such findings is unclear. On the other hand, it was reported that the retinal circadian clock synchronizes directly to light/dark cycles, which is independent of the SCN phase (59). Our results from the retina seem to agree with this finding, showing uncoupled phase relationships between the SCN and retina in mutants.

The miR-183/96/182 cluster is highly expressed in the neural system, particularly in sensory organs such as the inner ear and retina (39, 60). Knocking out of members of the *miR-183/96/182* cluster resulted in retinal degeneration (38, 41–43). The retina is not only a sensory organ but is also a self-sustained circadian clock (61). As the only source of photic input to the SCN in mammals, the retina has been proposed to contribute to the overall circadian organization (62). However, experiments from retinal dysfunction in rodents, generated either by a genetic approach or surgical approaches, had divergent observations. Most of them showed no effects on period length and phase while some of them found phase change or shortened period (63). It has been demonstrated that the *miR-183/96/182* cluster deficient mice have severely degenerated retinas (38, 39). However, in our study the retina of the mutant still exhibited sustained *Per2::Luc* rhythmicity, and normal pupillary light reflex mediated by melanopsin-containing retinal ganglion cells. The circadian mRNA profiling in the present study showed that the retina had dampened expression of core circadian genes in constant darkness but not in LD entrained condition. All of these suggest that the circadian changes may not be due to a significant contribution from retinal degeneration.

Intriguingly, the *miR-183/96/182* cluster deficient mice showed a significant circadian behavioral abnormality. The homozygous mutants became arrhythmic in constant darkness conditions. Although we observed various circadian alterations at both the cellular and tissue levels in different types of *miR-183/96/182* mutants, the cells and tissues still exhibited sustained rhythmicity. Behavior, however, does not necessarily reflect cell-autonomous clock phenotypes (64). The discrepancies between the circadian phenotypes at the behavioral and cellular levels may suggest that the *miR-183/96/182* cluster not only regulates circadian rhythms in a cell-autonomous way but may also affect the circadian system at a higher neuronal network level. Furthermore, uncoupled phase relationships between the SCN and lung in mutants may also suggest that something might be wrong in sending signals to the downstream or peripheral clocks from mutant SCN. Thus, despite confirmation of the cell-autonomous function of the miR-183/96/182 cluster, their functions in circadian neuronal circuits, including coupling, should not be excluded.

In summary, our study revealed that the miR-183/96/182 cluster has functional impacts on circadian rhythms in mammals. They can modulate circadian systems either by direct targeting of the core clock machinery or through a more indirect mechanisms that ultimately feeds into the circadian clock. Our genome-wide miRNA screen provides a valuable resource and insights into the new layer control of the circadian networks by non-coding RNAs.

## Materials and Methods

### Animals

All animal care and experimental procedures complied with University of Southern California guidelines for the care and use of animals and were approved by the University of Southern California Animal Care and Use Committee under protocol #20826.

The *miR-183/96/182* cluster gene trap mice (miR183C GT) were obtained as a gift from Dr. Changchun Xiao at The Scripps Research Institute. The *miR-183C^GT/GT^*; *Per2^Luc^* reporter mice were generated by crossing *miR-183C^GT/+^* mice with *Per2^Luc^* knock-in mice. All mice were housed in a temperature- and humidity-controlled room with food and water *ad libitum*. For time-course tissue dissections, mice of each genotype at 2–3 months of age were euthanized by isoflurane at each circadian time point, and the tissues were isolated, snap-frozen, and stored at - 80°C until processing for mRNA or protein analyses. For dissections of tissues during the dark phase, euthanasia of mice and dissections of eyes or retinas were performed under infrared illumination by a red lamp, and then the other tissues were dissected under the regular lights.

### Wheel-Running Activity

Mice were individually housed in cages equipped with running wheels, with food and water *ad libitum*. Mice were first entrained under 12 h/12 h light/dark conditions for 2 weeks, and then were released to constant darkness for 3 weeks. A Clock Lab system (Actimetrics) was used to collect wheel-running activity, and control the timing of light changes. LED lights were given as a square wave 12 h/12 h light/dark cycle with an irradiance of 5 W/m^2^ during the light phase.

### Total Activity

Locomotor activity was measured in polycarbonate cages (42 × 22 × 20 cm) placed into frames (25.5 × 47 cm) mounted with two levels of photocell beams at 2 and 7 cm above the bottom of the cage (San Diego Instruments, San Diego, CA, USA). These two sets of beams allowed for the recording of both horizontal (locomotion) and vertical (rearing) behaviors. The total activity was calculated as the sum of both types of behaviors. The mice were housed in the activity boxes with food and water *ad libitum*. They were first given 1 day to habituate to the testing environment. Immediately following this habituation, the mice were tested under a standard 12 h light / 12 h dark cycles for 7 days, and then constant darkness for 14 days.

### miRNA screen

*Per2-dLuc* and *Bmal1-dLuc* reporters cells (34) were reverse transfected with miRNAs in 384- well plates. Briefly, 20 µl of transfection reagent mixture (0.045 µl RNAiMax in Opti-MEM) was added to each well which was pre-spotted with miRNA (0.4 pmol). After 5 minutes of incubation at room temperature, 20 µl of cells (2500 cells/well) were added on top of the transfection mixture, and then were incubated at 37°C in 5% CO_2_ condition for 48 hours before recording. To record bioluminescence of the cells, the old medium was replaced with 60 µl HEPES-buffered explant medium supplemented with luciferin (1 mM) (GoldBio), 10% FBS and antibiotics. Plates were sealed with an optically clear film and were recorded in a plate reader (Synergy, BioTek) at a 2-hour interval for 5 days under 35°C condition. AllStars Negative Control siRNA, which does not target any gene in mammalian cells and therefore has no effects on circadian rhythms, was used as the negative control. An siRNA targeting luciferase was used as the luminescence signal control, and an siRNA targeting ubiquitous human cell survival genes was used as the cell survival control. Some miRNAs that were reported to modulate circadian rhythms (29–31, 65–67) were screened in the U2OS reporter cells for positive controls prior to the genome-wide miRNA screen. Among them, none of the miRNAs had significant and consistant effects on circadian period lengths, but only miRNA-142-3p robustly lowered the amplitude by targeting BMAL1 (29, 30). We therefore used miRNA-142-3p as a positive control for amplitude. CRY2 siRNA was used as another positive control for period length. The MultiCycle program (Actimetrics, Inc.) was used to analyze circadian bioluminescence data. Data was normalized (running average 24 hour methods) to detrend the baseline, and then the circadian parameters such as period length and amplitude were estimated using the best-fit sine wave analysis.

### CRISPR-Cas9 knockout of miRNAs in human cells

The knockout of miRNA was performed by the CRISPR-Cas9 system with paired synthetic crRNAs. Synthetic crRNAs were designed by the Dharmacon online CRISPR Design Tool. The high specificity scoring crRNAs which targeted sequences either in or close to the miRNA stem-loop were selected. Cells of a clonal U2OS *Per2-dLuc* cell line were seeded in a 96-well plate at a density of 10,000 cells per well one day prior to the transfection. Two pairs of 50nM crRNA:tracrRNA complex, and 200ng Cas9 (PuroR) plasmid which express a puromycin resistant gene were co-transfected by 0.4 μL/well DharmaFECT Duo reagent (GE Healthcare Dharmacon, Cat #T-2010-03). After 48 hours of transfection, the medium was replaced with complete medium containing 0.5 µg/mL puromycin, and cells were incubated for 3 days. Single-cell cloning was performed by serial dilution. The genotypes of single-cell clones were first determined by PCR and 2% agarose gel electrophoresis. The positive clones were further validated by Sanger sequencing. Sequences of crRNAs and PCR primers flanking the cleavage sites are listed in Table S5.

### Luciferase Assays

In a 384-well plate, the wells were pre-coated with 0.02% L-Lysine solutions, and then HEK-293T cells were plated at a density of 4000 cells/well. After 24 hours, the cells were co-transfected with 10ng 3’UTR reporter plasmid and 20 nM miRNA mimics per well. The transient transfections were performed using 0.16 µl of DharmaFECT DUO (Dharmacon) per well. Luciferase activity was measured 48 hours after transfection using the Bright-Glo™ Luciferase Assay System (Promega, Cat #E2610) on a microplate reader (Infinite M200, Tecan) according to the manufacturer’s instructions.

### RNA Isolation

Total RNA was extracted using the RNAzol method combined with columns. Briefly, cells or tissues were homogenized in 500 µl RNAzol RT (MRC), and mixed with 200 µl of RNase free water. The mixture was then vortexed for 15□seconds and incubated for 15□minutes at room temperature, followed by 15□minutes centrifugation at 16 000□g. The supernatant was transferred to a fresh tube with 8□μl of 4-Bromoanisol and vortexed for 15□seconds. After incubation at room temperature for 5□minutes, the solution was centrifuged for 10□minutes at 12 000□g. The supernatant contains RNA and was transferred to a fresh tube. The mixture of the RNA containing supernantant and 1 volume of isopropanol was filtered by Zymo-Spin™ ICG Column (Zymo). RNA was washed twice with Direct-zol™ RNA PreWash buffer, and once with RNA Wash Buffer on column, and then elute with 10-20 µl of RNase free water. For large size tissues such as eye balls and lung, the reagent volumes doubled accordingly.

### Quantitative RT-PCR

A two-tailed RT-PCR method (68) was used for miRNA quantification. Reverse transcription (RT) reactions were performed using the iScript select cDNA kit (BioRad) in a total reaction volume of 10 µl, which contained 1x reaction buffer, 1 µl of GSP enhancer solution, mix of 0.5 µM Two-tailed RT primers, 0.5 µl reverse transcriptase and 100 ng total RNA. The reactions were incubated at 42°C for 45 minutes, and 85°C for 5 minutes in a thermocycler. Reverse transcription (RT) reactions for mRNA were performed in a total reaction volume of 10 µl, which contained 1x SuperScript IV VILO Master Mix (Invitrogen) and 200 ng RNA. The reactions were incubated at 25°C for 10 min, 50°C for 10 minutes and 85°C for 5 minutes in a thermocycler.

Methods for qPCR were the same for both miRNA and mRNA. One total volume of 10 µl reaction contained 1x Maxima SYBR Green qPCR Master Mix (Thermo Scientific), 0.5 µM forward and reverse primers, and 3 µl of diluted cDNA template. qPCR was performed in a CFX 384 Real Time Detection System (Bio-Rad) at 95°C for 10 minutes, 40 cycles of 95°C for 15 seconds, and 60°C for 60 seconds, followed by melting-curve analysis. The primers used in qPCR analysis are listed in Table S6.

### Western Blotting

Whole-cell extracts were prepared using lysis buffer containing 50 mM Tris (pH8), 1% TX-100, 150 mM NaCl, 12 mM Sodium Dexcycolate, 0.1% SDS, 10 mM EDTA, and protease inhibitor. After incubation on ice for 10 minutes, samples were centrifuged at 16 000 g, at 4°C for 20 minutes. Proteins were quantified by DC Protein Assay Kit (BioRad). Gel electrophoresis was run in Criterion TGX Stain-Free Gels (BioRad). Antibodies were anti-PER2 polyclonal (ProteinTech # 20359-1-AP) and anti-GAPDH (ProteinTech # 10494-1-AP).

### Bioluminescence recording of tissues from *Per2^Luc^* reporter mice

The *miR-183C^GT/GT^; Per2^Luc^* mice and littermate wild-type control mice were euthanized by isoflurane 2-3 hours before lights off (ZT 9–ZT10) and tissues were immediately removed into ice-cold HBSS (Gibco). Coronal brain slices containing SCN (400-µm thickness) were cut using a tissue slicer (Stoelting). The bilateral SCN tissue without optic chiasm attached was further dissected out from the brain slice with a sterile scalpel under a dissecting microscope. The SCN cultures were cultured on cell culture inserts (PICMORG50; Millipore) in sealed 35-mm culture dishes containing 1.5mL DMEM supplemented with 1 x B-27 plus (Gibco), 352.5 μg/mL sodium bicarbonate, 10 mM Hepes (Gibco), 25 U/mL penicillin, 25 μg/mL streptomycin (Gibco), and 0.1 mM D-luciferin potassium salt (GoldBio). Retinas were isolated from the eye balls and then gently placed on culture inserts (PICMORG50; Millipore) with ganglion cell down in Neural basal plus medium (Gibco) containing 1 x B-27 plus (Gibco) and 2% FBS (Atlanta Biologicals), 25 U/mL penicillin, 25 μg/mL streptomycin (Gibco), and 2 mM GlutaMAX (Gibco). Bioluminescence was measured with a LumiCycle instrument (Actimetrics, Wilmette, IL). Cultures were maintained in a light-tight incubator at 37°C for one week. Data was analyzed by using the LM fit (damped sin) method in LumiCycle data analysis software (Actimetrics). The time of the peak of luminescence on the first day of constant conditions (first day in LumiCycle) was determined as the time of peak of the sine wave on that day.

### Statistical Analysis

The statistical significance was determined by unpaired Student’s two tailed t-test if there were only two groups of data to compare. In cases of comparing multiple conditions, ordinary one-way ANOVA and Tukey post hoc tests were applied. P values of less than 0.05 were considered significant.

## Acknowledgments

We thank Dr. Chaungchun Xiao at The Scripps Research Institute for miR-183/96/182 cluster GT mice, and Dr. Amanda Roberts at The Scripps Research Institute Animal Models Core Facility for assistance of non-wheel behavioral assessment. We thank Dr. Tsuyoshi Hirota at Nagoya University for giving comments on the manuscript. We also thank our colleagues from the Kay lab for discussion and suggestions. This work was supported by National Institute of Diabetes and Digestive and Kidney Diseases Grant 5R01DK108087 to S.A.K.

## Supplementary Information

**Fig. S1.**
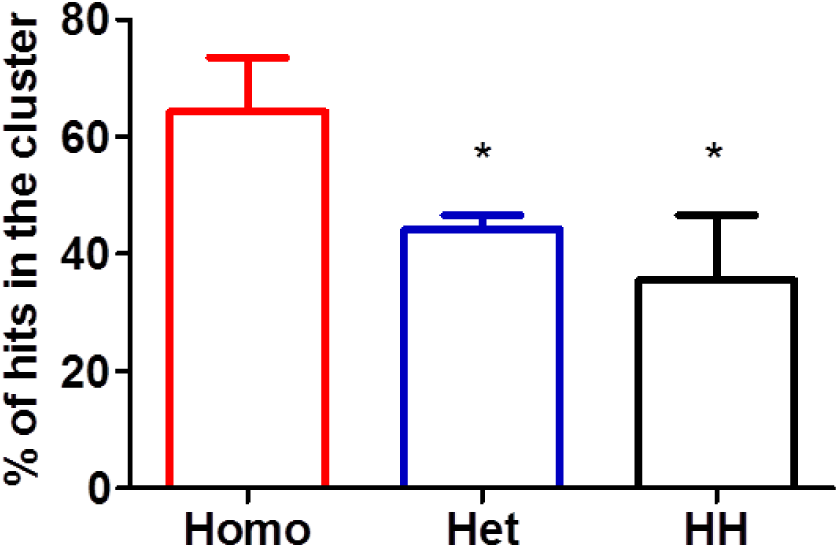
Homo-seed clusters (Homo) have a significantly higher percentage of candidate miRNA enrichment than that of hetero-seed clusters (Het) and homo-hetero-seed clusters (HH). * *p* < 0.05. Data represent the mean ± SEM.

**Fig. S2.**
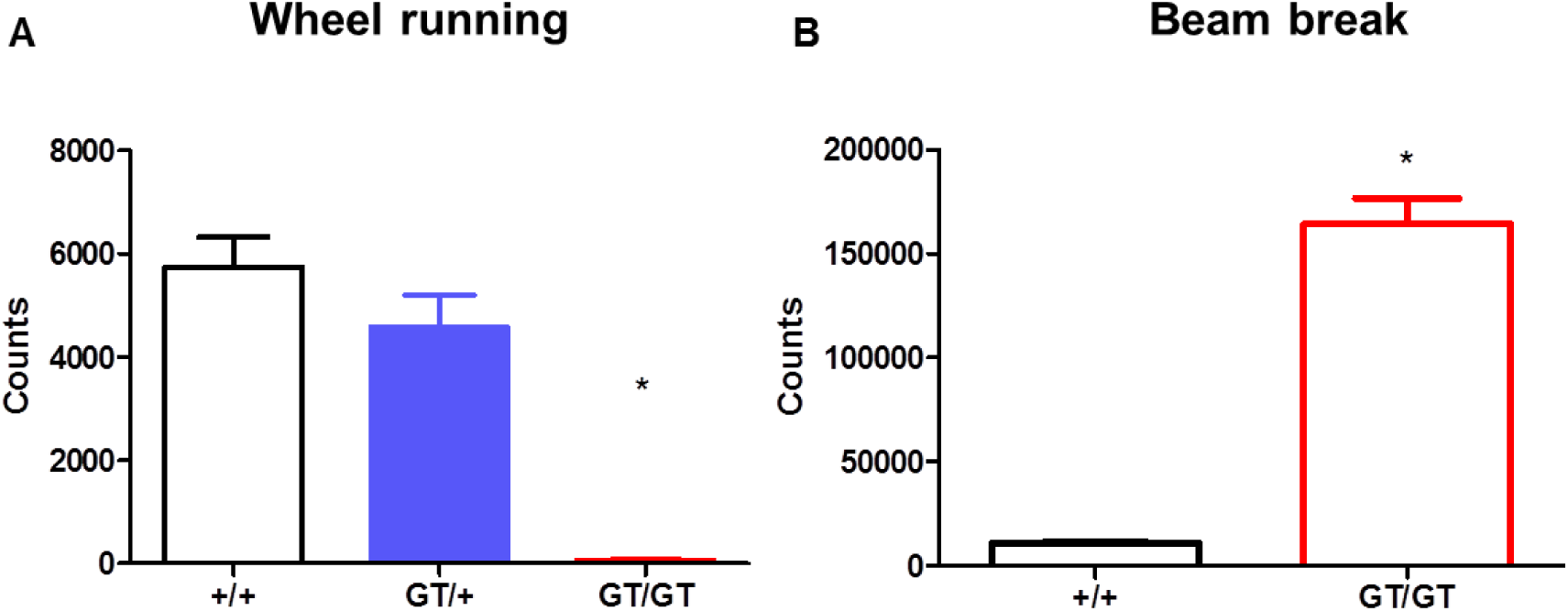
(A) *miR-183C^GT/GT^* mice had significantly lower daily activity on running wheels than wild-type mice. (B) The general daily activity of *miR-183C^GT/GT^* mice was significantly higher than that of wild-type mice. Data represent the mean ± SEM.

**Fig. S3.**
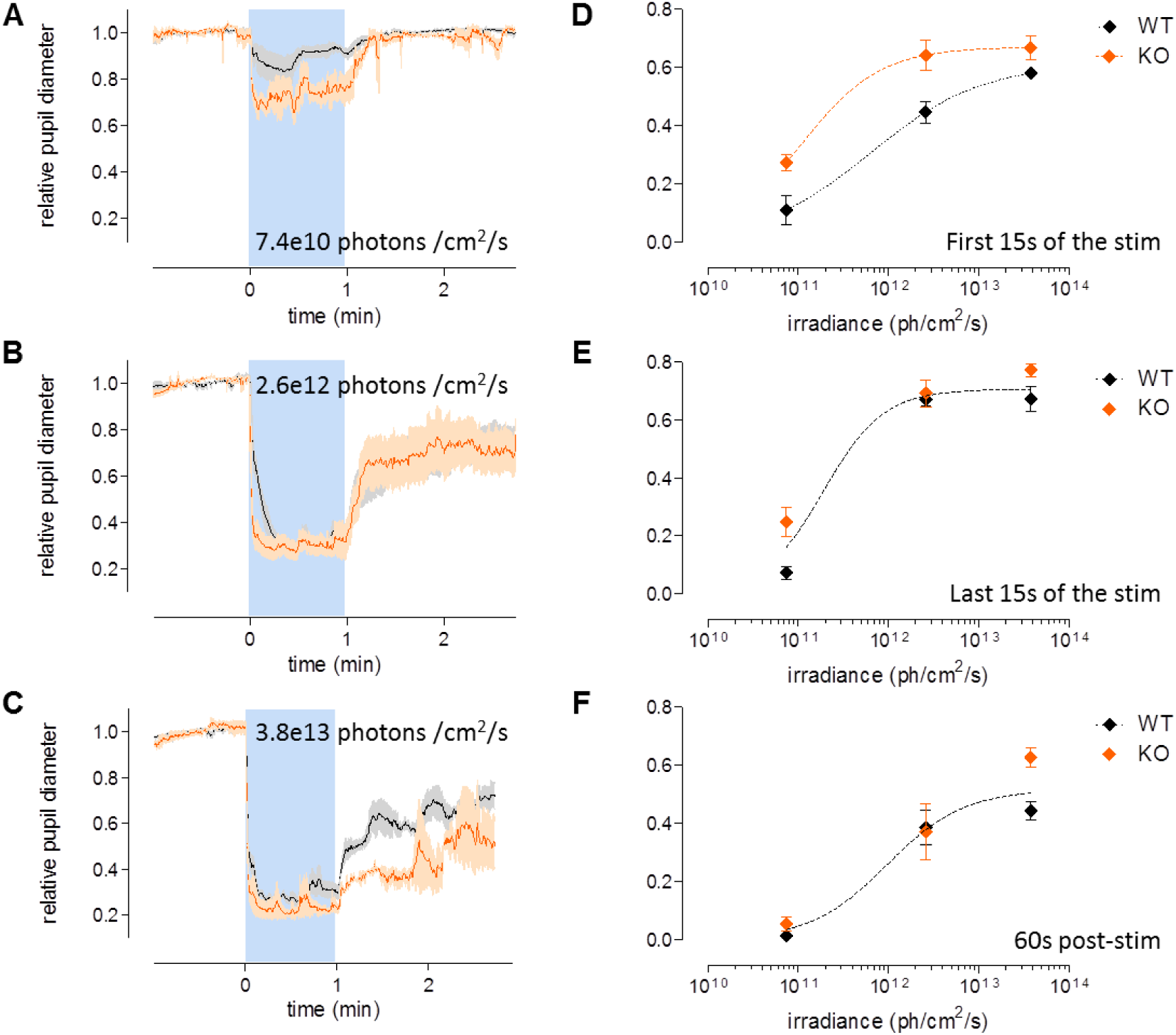
Pupillary light reflex mediated by melanopsin was not diminished. (A-C) Average pupil diameter (normalized to baseline) in response to 1 min monochromatic blue light stimulation at increasing irradiance (480nm, 7.4×10^10^, 2.6×10^12^, and 3.8×10^13^ photons/cm2/s). Each 5-min recording sequence consisted of 1 min of darkness for baseline, 1 min of monochromatic 480 nm light for stimulation (blue shade), and finally 3 min of darkness for recovery. Results from the first 15 seconds during stimulation, the last 15 seconds during stimulation, and the first 60 seconds of post-stimulation were used for assessing the responses between mice of different genotypes, and were summarized in (D-F). Results from the 60s post-stimulation of 2.6×10^12^ photons indicated that the pupillary light reflex mediated by melanopsin were comparable between *miR-183C^GT/GT^* (WT) and *miR-183C^GT/GT^* (KO) mice. Data represent the mean ± SEM.

**Fig. S4.**
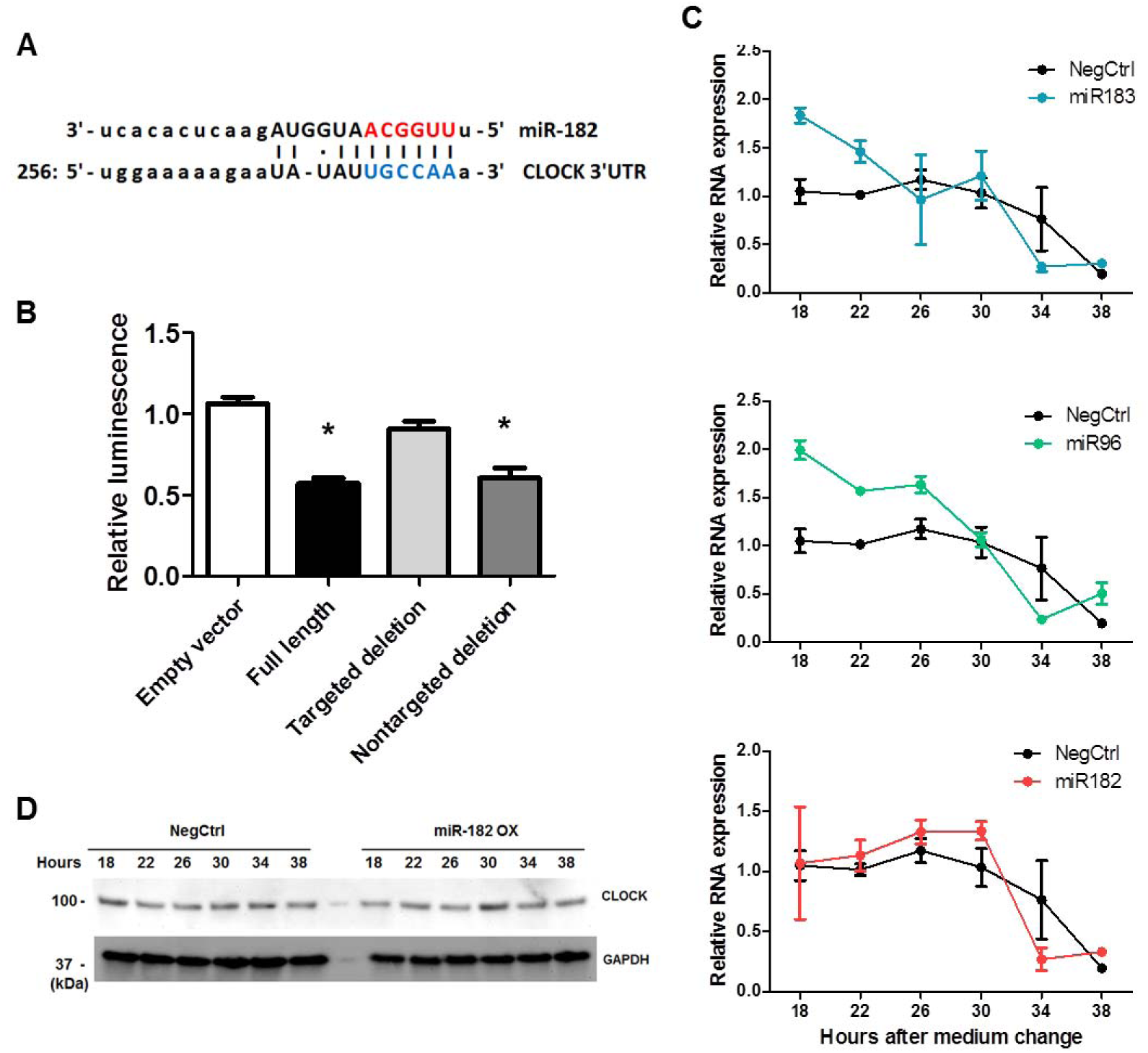
(A) Sequence alignment between miR-182 and its putative binding site (blue letters) in CLOCK-3’UTR. Red letters indicate canonical seed nucleotides. (B) HEK-293T cells were transfected with a luciferase reporter plasmid containing either no 3’UTR, or the CLOCK-3’UTR with wild-type, mutated target site, or mutated non-target site of miR-182. (C) *CLOCK* mRNA expression in U2OS cells transfected with miRNA mimics of each member of the miR-183/96/182 cluster. RNA was sampled every 4 hours 18 hours after medium change. (D) U2OS cells were transfected with either control or miR-182 mimics. CLOCK protein levels were analyzed by western blot 18 hours after medium change. Data represent the mean ± SEM.

**Fig. S5.**
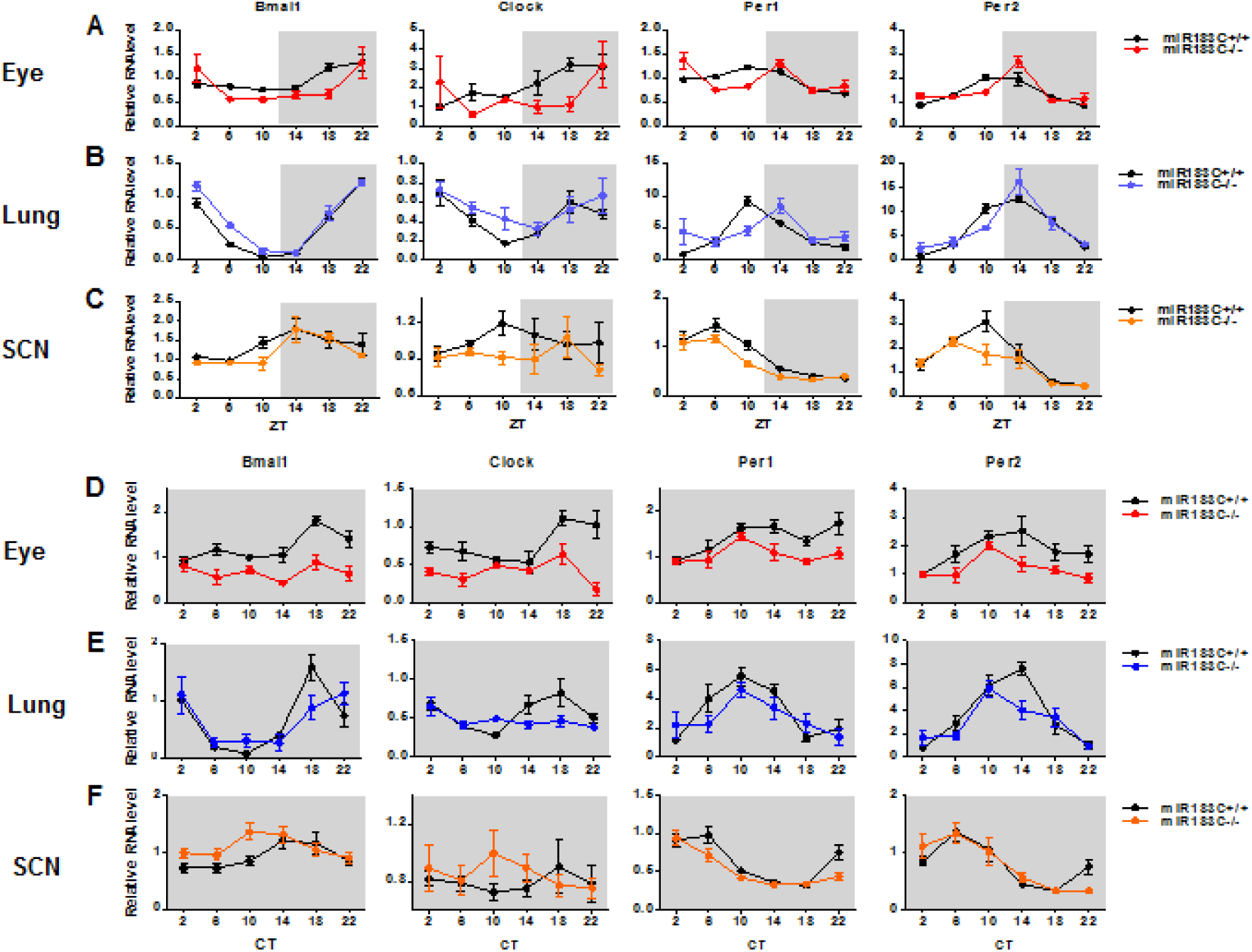
Tissues of the whole eye ball (A and D), the lung (B and E), and the SCN (C and F) from either *miR-183C^+/+^* mice or *miR-183C^GT/GT^* mice were sampled every 4 hours across 24 hours either under LD (top panel) or DD (bottom panel) conditions. The relative mRNA of core circadian genes was measured by qPCR (n = 4∼5). Data represent the mean ± SEM.

**Table S1.**
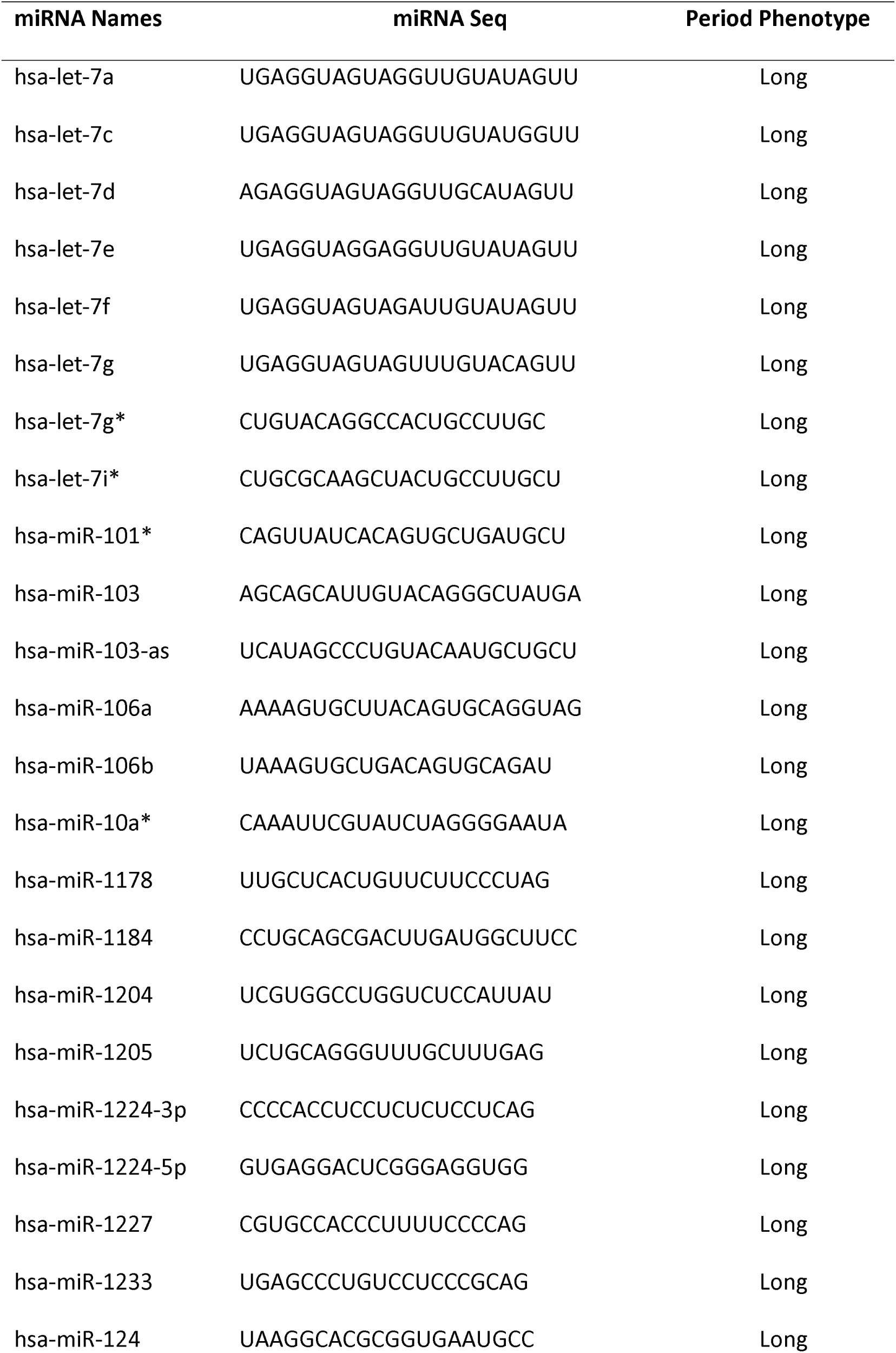
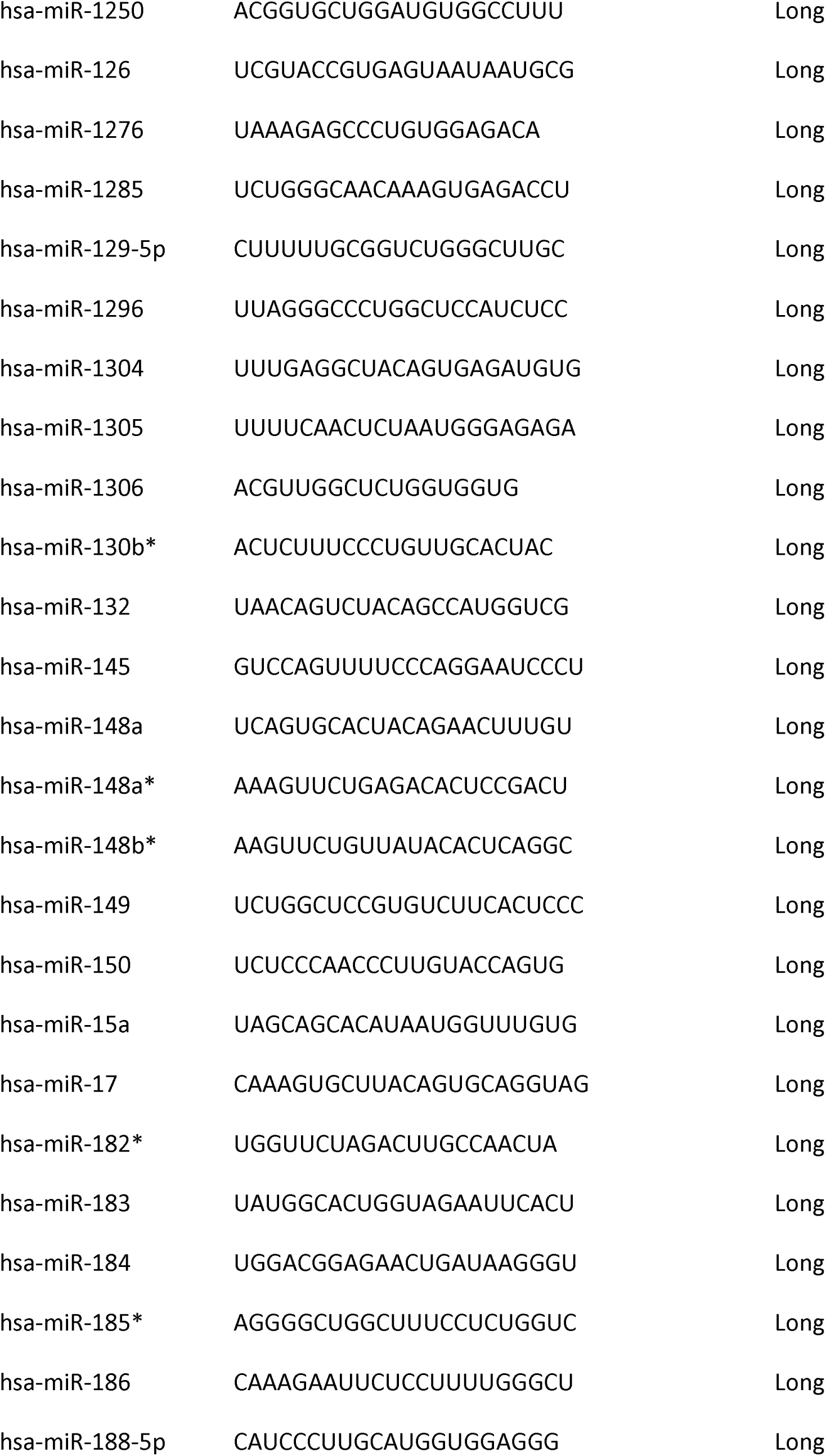
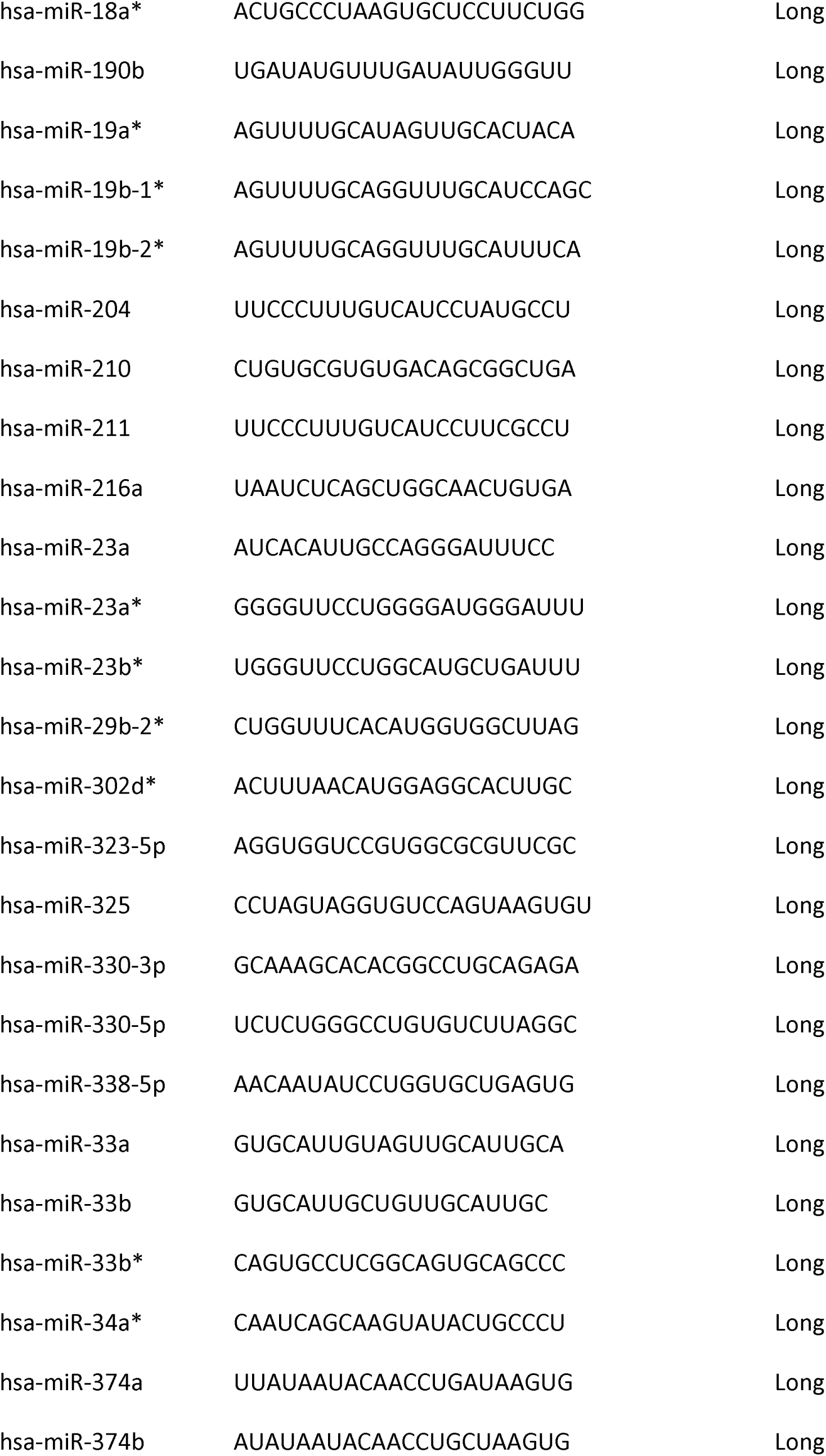
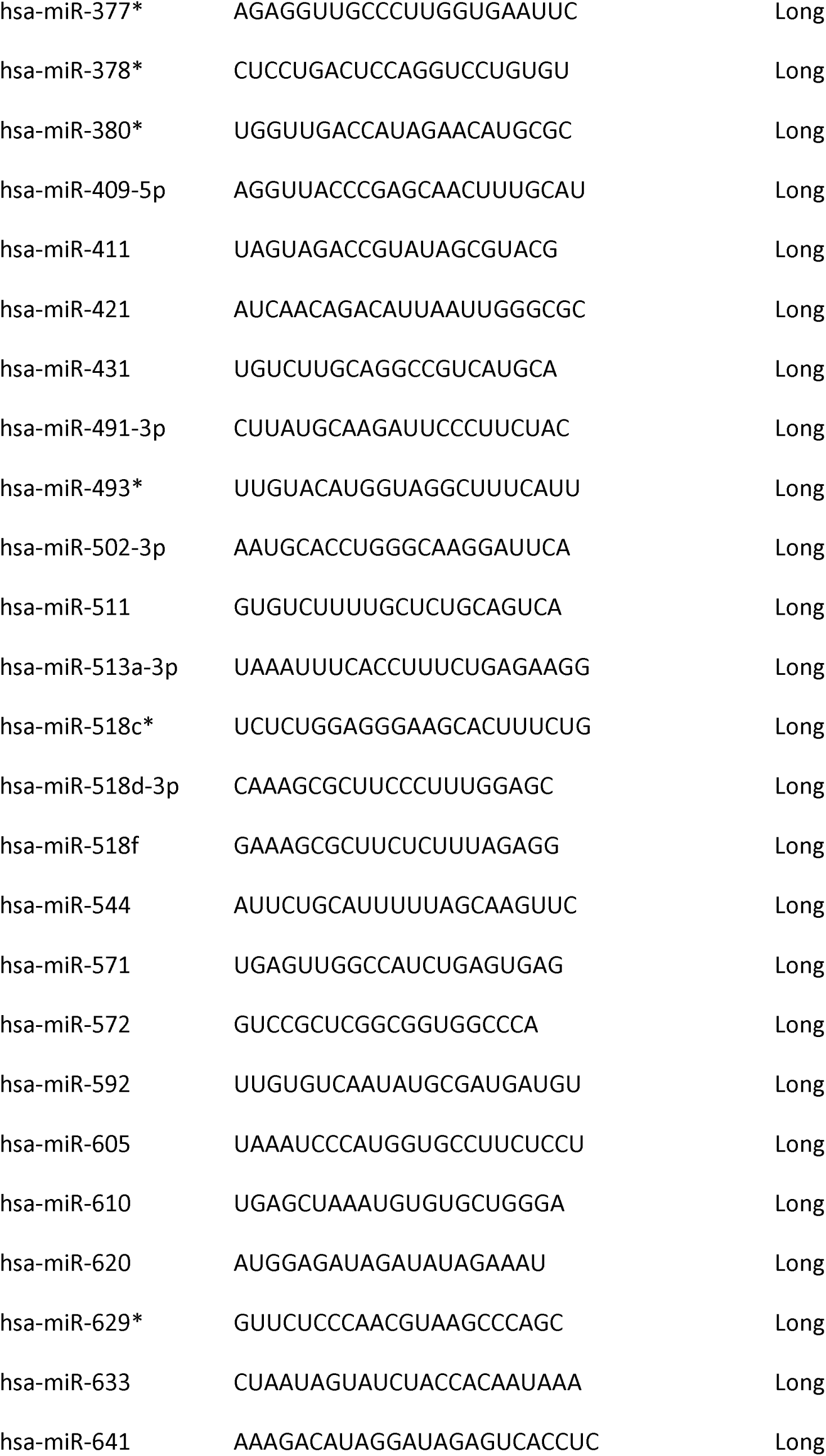
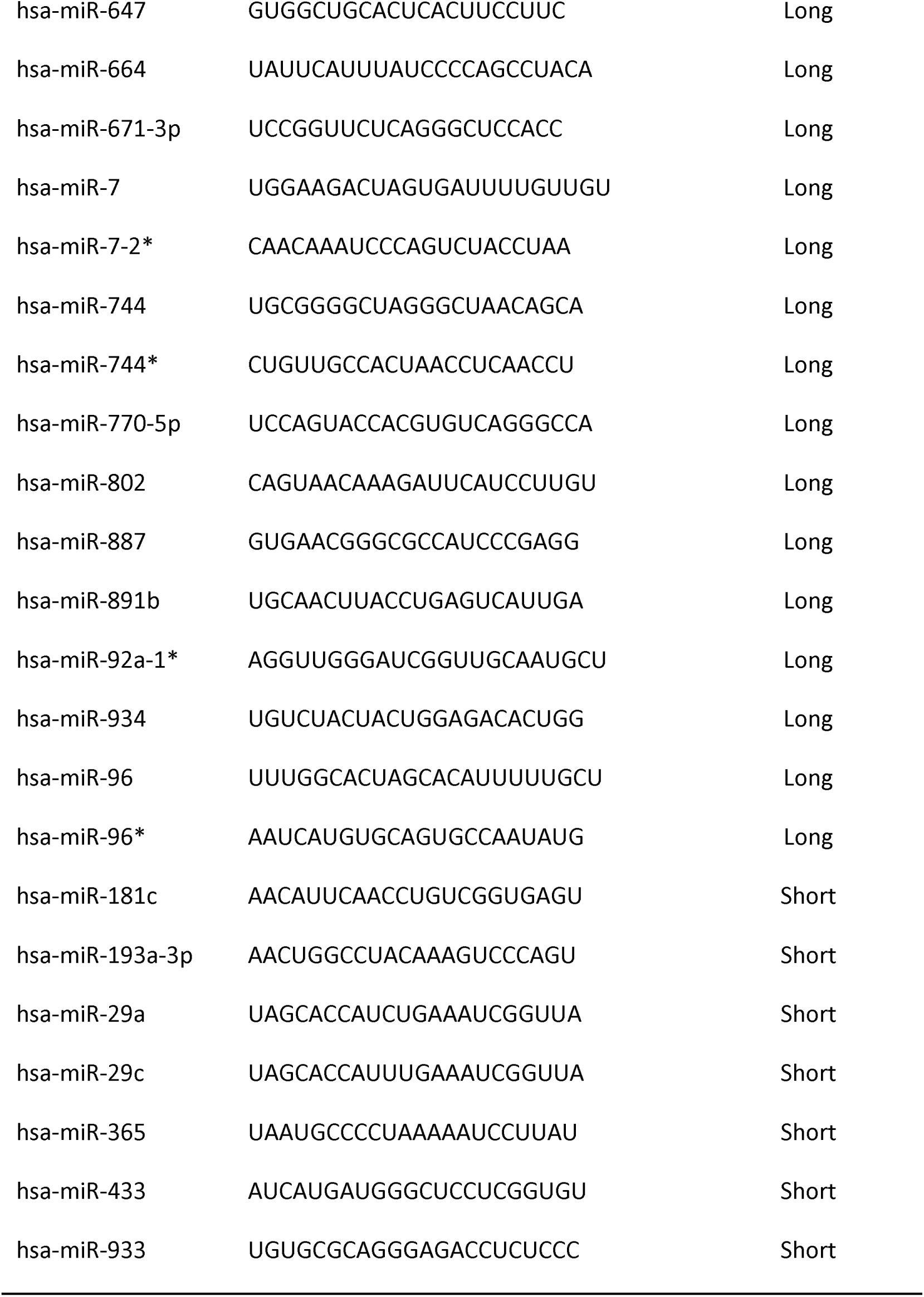
Candidate miRNA hits affecting period length.

**Table S2.** Gene family information for each candidate miRNA hit.

| **miRNA Names** | **Gene family** |
| --- | --- |
| hsa-let-7f | let-7 |
| hsa-let-7a | let-7 |
| hsa-let-7d | let-7 |
| hsa-let-7c | let-7 |
| hsa-let-7e | let-7 |
| hsa-let-7g* | let-7 |
| hsa-let-7g | let-7 |
| hsa-let-7i* | let-7 |
| hsa-miR-10a* | mir-10 |
| hsa-miR-101* | mir-101 |
| hsa-miR-103-as | mir-103 |
| hsa-miR-103 | mir-103 |
| hsa-miR-1178 | mir-1178 |
| hsa-miR-1184 | mir-1184 |
| hsa-miR-1204 | mir-1204 |
| hsa-miR-1205 | mir-1205 |
| hsa-miR-1224-5p | mir-1224 |
| hsa-miR-1224-3p | mir-1224 |
| hsa-miR-1227 | mir-1227 |
| hsa-miR-1233 | mir-1233 |
| hsa-miR-124 | mir-124 |
| hsa-miR-1250 | mir-1250 |
| hsa-miR-126 | mir-126 |
| hsa-miR-1276 | mir-1276 |
| hsa-miR-1285 | mir-1285 |
| hsa-miR-129-5p | mir-129 |
| hsa-miR-1296 | mir-1296 |
| hsa-miR-130b* | mir-130 |
| hsa-miR-1304 | mir-1304 |
| hsa-miR-1305 | mir-1305 |
| hsa-miR-1306 | mir-1306 |
| hsa-miR-132 | mir-132 |
| hsa-miR-145 | mir-145 |
| hsa-miR-148b* | mir-148 |
| hsa-miR-148a | mir-148 |
| hsa-miR-148a* | mir-148 |
| hsa-miR-149 | mir-149 |
| hsa-miR-15a | mir-15 |
| hsa-miR-150 | mir-150 |
| hsa-miR-323-5p | mir-154 |
| hsa-miR-377* | mir-154 |
| hsa-miR-409-5p | mir-154 |
| hsa-miR-106a | mir-17 |
| hsa-miR-106b | mir-17 |
| hsa-miR-18a* | mir-17 |
| hsa-miR-17 | mir-17 |
| hsa-miR-181c | mir-181 |
| hsa-miR-182* | mir-182 |
| hsa-miR-183 | mir-183 |
| hsa-miR-184 | mir-184 |
| hsa-miR-185* | mir-185 |
| hsa-miR-186 | mir-186 |
| hsa-miR-188-5p | mir-188 |
| hsa-miR-19b-2* | mir-19 |
| hsa-miR-19b-1* | mir-19 |
| hsa-miR-19a* | mir-19 |
| hsa-miR-190b | mir-190 |
| hsa-miR-193a-3p | mir-193 |
| hsa-miR-204 | mir-204 |
| hsa-miR-211 | mir-204 |
| hsa-miR-210 | mir-210 |
| hsa-miR-216a | mir-216 |
| hsa-miR-23b* | mir-23 |
| hsa-miR-23a* | mir-23 |
| hsa-miR-23a | mir-23 |
| hsa-miR-92a-1* | mir-25 |
| hsa-miR-29a | mir-29 |
| hsa-miR-29c | mir-29 |
| hsa-miR-29b-2* | mir-29 |
| hsa-miR-302d* | mir-302 |
| hsa-miR-325 | mir-325 |
| hsa-miR-33b* | mir-33 |
| hsa-miR-33b | mir-33 |
| hsa-miR-33a | mir-33 |
| hsa-miR-330-3p | mir-330 |
| hsa-miR-330-5p | mir-330 |
| hsa-miR-338-5p | mir-338 |
| hsa-miR-34a* | mir-34 |
| hsa-miR-365 | mir-365 |
| hsa-miR-374a | mir-374 |
| hsa-miR-374b | mir-374 |
| hsa-miR-378* | mir-378 |
| hsa-miR-380* | mir-379 |
| hsa-miR-411 | mir-379 |
| hsa-miR-431 | mir-431 |
| hsa-miR-433 | mir-433 |
| hsa-miR-491-3p | mir-491 |
| hsa-miR-493* | mir-493 |
| hsa-miR-502-3p | mir-500 |
| hsa-miR-513a-3p | mir-506 |
| hsa-miR-511 | mir-506 |
| hsa-miR-518c* | mir-515 |
| hsa-miR-518f | mir-515 |
| hsa-miR-518d-3p | mir-515 |
| hsa-miR-544 | mir-544 |
| hsa-miR-571 | mir-571 |
| hsa-miR-572 | mir-572 |
| hsa-miR-592 | mir-592 |
| hsa-miR-605 | mir-605 |
| hsa-miR-610 | mir-610 |
| hsa-miR-629* | mir-629 |
| hsa-miR-633 | mir-633 |
| hsa-miR-641 | mir-641 |
| hsa-miR-647 | mir-647 |
| hsa-miR-664 | mir-664 |
| hsa-miR-671-3p | mir-671 |
| hsa-miR-7-2* | mir-7 |
| hsa-miR-7 | mir-7 |
| hsa-miR-744* | mir-744 |
| hsa-miR-744 | mir-744 |
| hsa-miR-770-5p | mir-770 |
| hsa-miR-802 | mir-802 |
| hsa-miR-887 | mir-887 |
| hsa-miR-891b | mir-891 |
| hsa-miR-933 | mir-933 |
| hsa-miR-934 | mir-934 |
| hsa-miR-421 | mir-95 |
| hsa-miR-96* | mir-96 |
| hsa-miR-96 | mir-96 |

**Table S3.** Gene cluster information for each candidate miRNA hit and the categories based on seed sequence similarity. Red color denotes the miRNAs identified from the screen.

| Category | miRNA cluster |
| --- | --- |
| Homo | let-7a-1; let-7f-1; let-7d |
| Homo | mir-29c; mir-29b-2 |
| Homo | mir-132; mir-212 |
| Homo | mir-16-1; mir-15a |
| Homo | mir-181c; mir-181d |
| Homo | mir-29a; mir-29b-1 |
| Homo | mir-301b; mir-130b |
| HH | mir-379; mir-411; mir-299; mir-380; mir-1197; mir-323a; mir-758; mir-329-1; mir-329-2; mir-494; mir-1193; mir-543; mir-495; mir-376c; mir-376a-2; mir-654; mir-376b; mir-376a-1; mir-300; mir-1185-1; mir-1185-2; mir-381; mir-487b; mir-539; mir-889; mir-544a; mir-655; mir-487a; mir-382; mir-134; mir-668; mir-485; mir-323b; mir-154; mir-496; mir-377; mir-541; mir-409; mir-412; mir-369; mir-410; mir-656 |
| HH | mir-17; mir-18a; mir-19a; mir-20a; mir-19b-1; mir-92a-1 |
| HH | 512-1; 512-2; 1323; 498; 520e; 515-1; 519e; 520f; 515-2; 519c; 1283-1; 520a; 526b; 519b; 525; 523; 518f; 520b; 518b; 526a-1; 520c; 518c; 524; 517a; 519d; 521-2; 520d; 517b; 520g; 516b-2; 526a-2; 518e; 518a-1; 518d; 516b-1; 518a-2; 517c; 520h; 521-1; 522; 519a-1; 527; 516a-1; 1283-2; 516a-2; 519a-2 |
| HH | mir-106a; mir-18b; mir-20b; mir-19b-2; mir-92a-2; mir-363 |
| HH | mir-106b; mir-93; mir-25 |
| HH | mir-302b; mir-302c; mir-302a; mir-302d; mir-367 |
| HH | mir-532; mir-188; mir-500a; mir-362; mir-501; mir-500b; mir-660; mir-502 |
| HH | mir-891b; mir-892b; mir-892a; mir-888; mir-890; mir-892c |
| HH | mir-183; mir-96; mir-182 |
| Hetero | mir-1250; mir-338; mir-657 |
| Hetero | mir-439; mir-337; mir-665; mir-431; mir-433; mir-127; mir-432; mir-136 |
| Hetero | mir-101; mir-3671 |
| Hetero | mir-1179; mir-7-2 |
| Hetero | mir-143; mir-145 |
| Hetero | mir-193b; mir-365a |
| Hetero | mir-217; mir-216a |
| Hetero | mir-23b; mir-27b; mir-24-1 |
| Hetero | mir-24-2; mir-27a; mir-23a |
| Hetero | mir-33b; mir-6777 |
| Hetero | mir-3618; mir-1306 |
| Hetero | mir-374a; mir-545 |
| Hetero | mir-374b; mir-421 |
| Hetero | mir-493; mir-337; mir-665 |
| Hetero | mir-507; mir-506; mir-513a-2 |
| Hetero | mir-647; mir-1914 |
| Hetero | mir-6789; mir-1227 |
| Hetero | mir-99a; let-7c; mir-125b-2 |
| Hetero | mir-99b; let-7e; mir-125a |

**Table S4.** Predicted targets for members of miR-183 cluster by different methods.

|  | **miR-183** | **miR-96** | **miR182** |
| --- | --- | --- | --- |
| BMAL1 | - | - | - |
| CLOCK | Diana | miRanda, Diana | miRanda, Diana |
| CRY1 | - | - | - |
| CRY2 | - | - | - |
| PER1 | - | - | - |
| PER2 | - | miRanda, Diana | Diana |

**Table S5.**
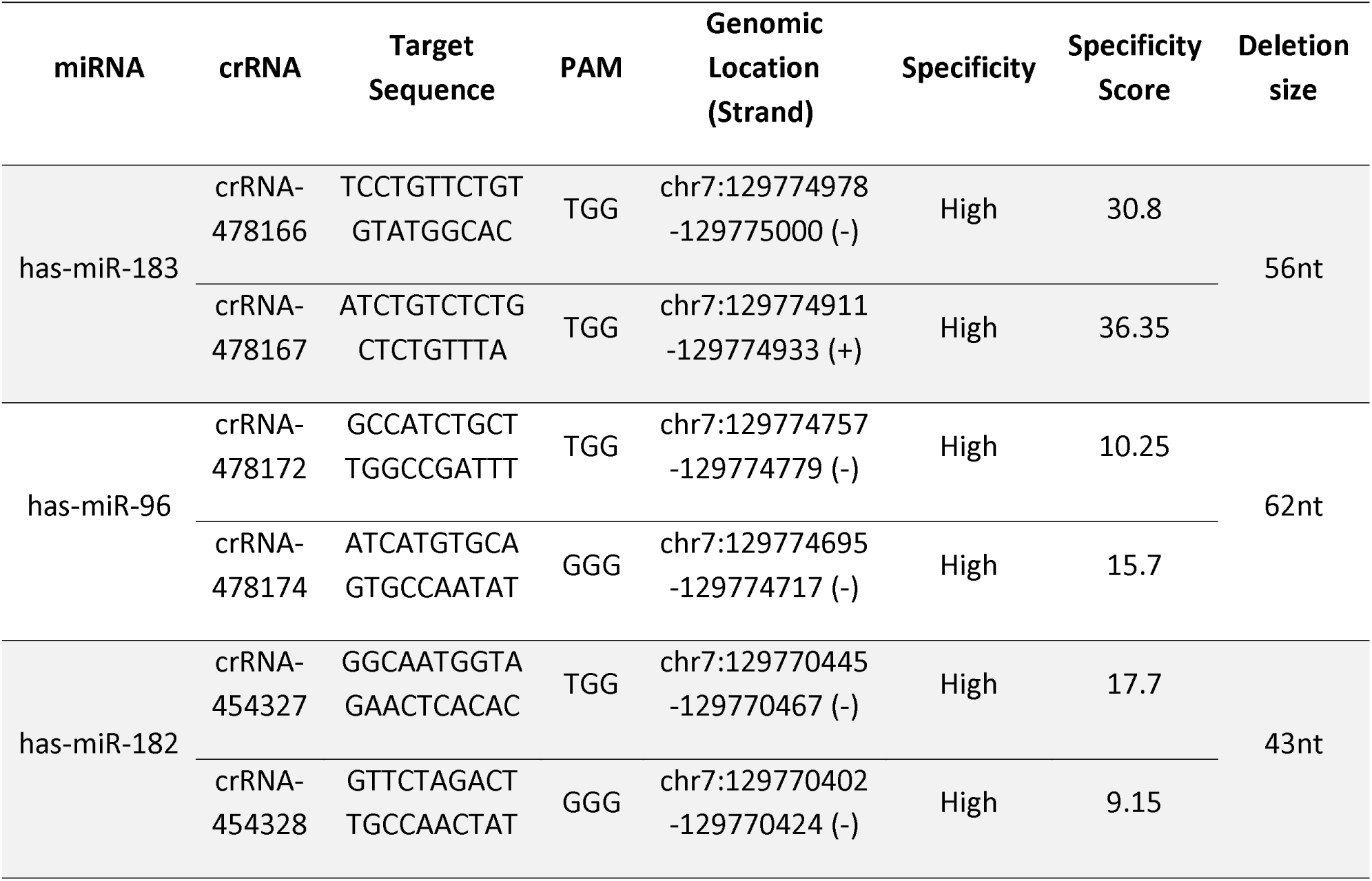
RNA sequences for CRISPR.

**Table S6.**
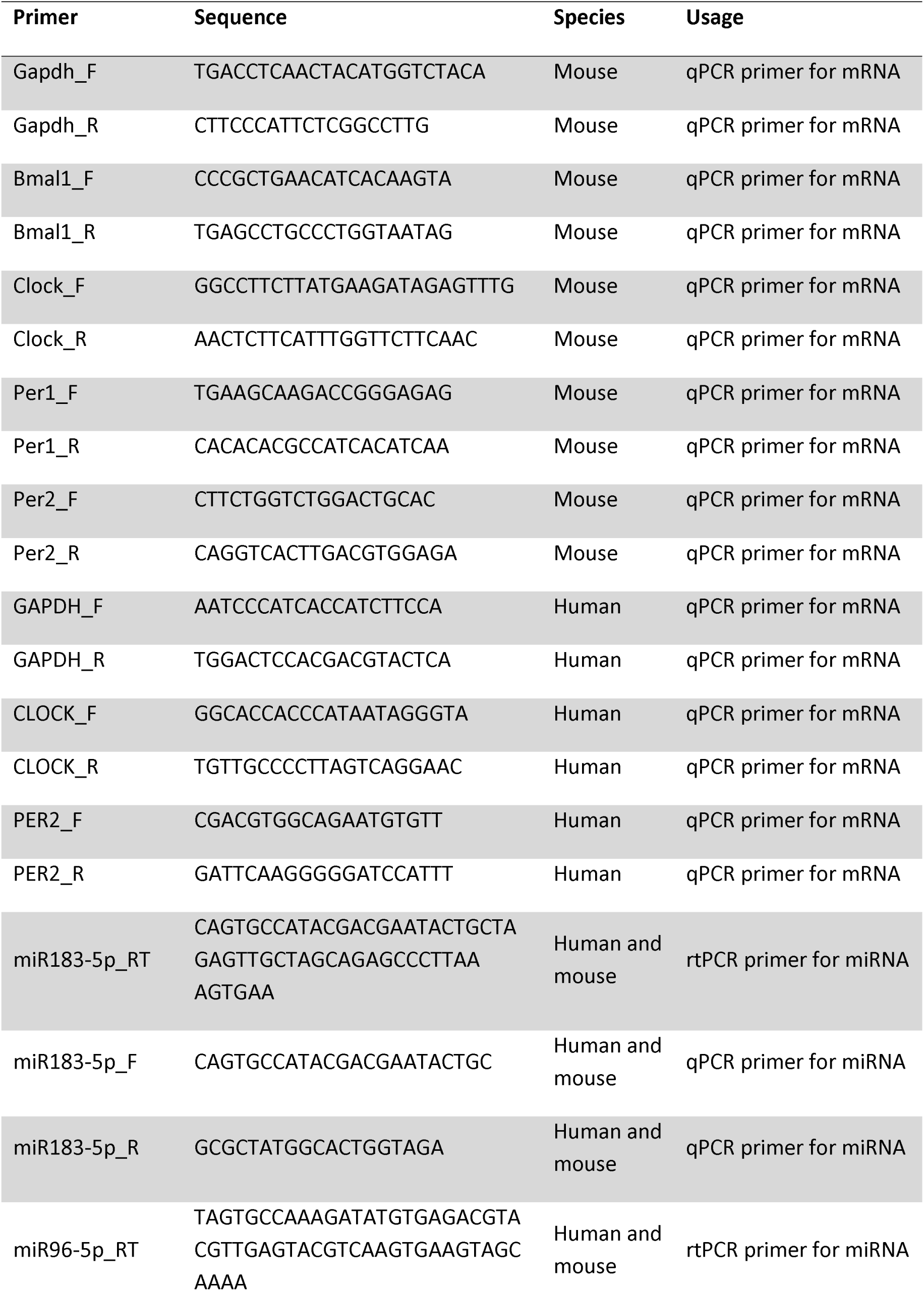
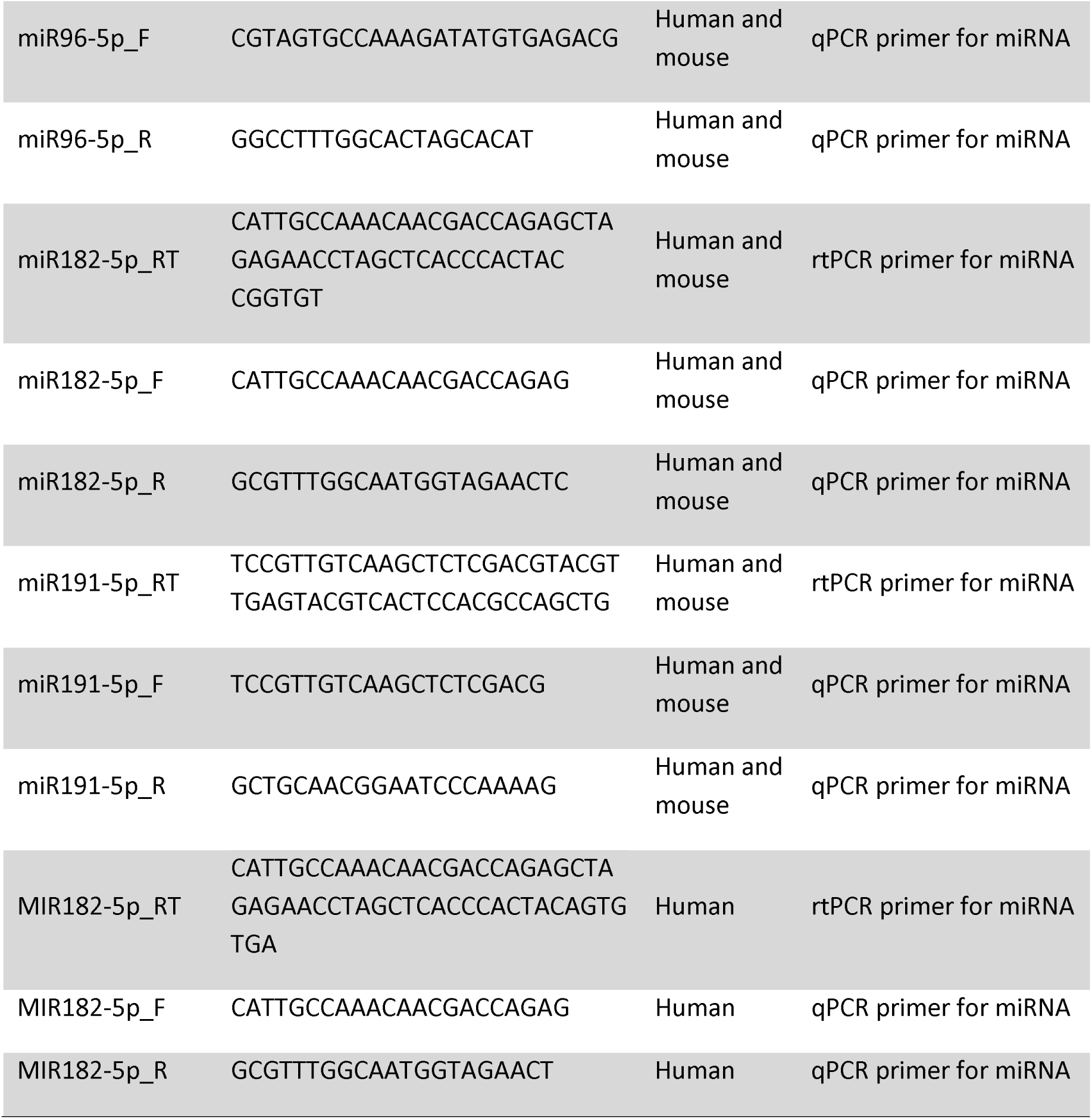
Primers for qPCR.

